# Antifungal potency and modes of action of a novel olive tree defensin against closely related ascomycete fungal pathogens

**DOI:** 10.1101/704072

**Authors:** Hui Li, Siva L. S. Velivelli, Dilip M. Shah

## Abstract

Antimicrobial peptides play a pivotal role in the innate immunity of plants. Defensins are cysteine-rich antifungal peptides with multiple mechanisms of action (MOA). A novel *Oleaceae*-specific defensin gene family was discovered in the genome sequences of the wild and cultivated species of a perennial olive tree, *Olea europaea*. Antifungal properties of an olive tree defensin OefDef1.1 were investigated against a necrotrophic ascomycete fungal pathogen *Botrytis cinerea in vitro* and *in planta*. OefDef1.1 displayed potent antifungal activity against this pathogen by rapidly permeabilizing the plasma membrane of the conidial and germling cells. Interestingly, it was translocated to the cytoplasm and induced reactive oxygen species in the germlings, but not in the conidia. In medium containing high concentrations of Na^1+^, antifungal activity of OefDef1.1 against *B. cinerea* was significantly reduced. In contrast, OefDef1.1_V1 variant in which the γ-core motif of OefDef1.1 was replaced by that of a *Medicago truncatula* defensin MtDef4 displayed Na^1+^-tolerant antifungal activity and was more potent in reducing the virulence of *B. cinerea in planta*. OefDef1.1 also exhibited potent antifungal activity against three hemibiotrophic ascomycete pathogens *Fusarium graminearum, F. oxysporum* and *F. virguliforme*. Significant differences were observed among the four pathogens in their responses to OefDef1.1 in growth medium with or without the high concentrations of Na^1+^. The varied responses of closely related ascomycete pathogens to this defensin have implications for engineering disease resistance in plants.

## Introduction

Antimicrobial peptides (AMPs) are recognized as important mediators of innate immunity in the plant kingdom providing first-line of defense against fungal and oomycete pathogens (van der Weerden *et al.*, 2013; Goyal & Mattoo, 2014). Among several classes of AMPs expressed in plants, small cysteine-rich defensins have been extensively studied for their antimicrobial properties, mechanisms of action (MOA) and ability to provide protection from fungal and oomycete pathogens in crops (Kaur *et al.*, 2011; de Coninck *et al.*, 2013; Lacerda *et al*., 2014; Cools *et al.*, 2017; Parisi *et al.*, 2018). There are now *c.* 1200 plant defensin sequences in the database (Shafee *et al.*, 2016). A vast majority of these peptides still remain to be studied at a functional level. Plant defensins are 45-54 amino acids in length and contain an invariant tetradisulfide array which confers stability to their pseudo-cyclic backbone. Because of their conserved cysteine signature, plant defensins share a similar 3D structure comprising a triple-stranded β-sheet and one α-helix, suggesting they evolved from a common ancestor (Van der Weerden & Anderson, 2013). Despite their structural similarity, plant defensins are highly varied in their primary amino acid sequences.

The best known property of cationic plant defensins is their ability to inhibit the growth of fungal pathogens *in vitro* and *in planta*. However, the MOA of only a small number of plant defensins have been studied in detail (Cools *et al.*, 2017; Parisi *et al.*, 2019). It is now recognized that plant defensins act using different MOA, but ultimately cause membrane disruption. Some plant defensins act extracellularly on fungi and bind to specific cell wall/plasma membrane resident sphingolipids, disrupt membrane integrity and activate cellular toxicity pathways. For example, the radish defensin RsAFP2 binds to glucosylceramide and this interaction results in the induction of cell wall stress, accumulation of ceramides and reactive oxygen species (ROS), and ultimately cell death (Thevissen *et al.*, 2012). During the last few years, several plant defensins that gain entry into fungal cells and target plasma membrane resident phospholipids have been characterized (Islam *et al.*, 2017; Parisi *et al.*, 2019). Plant defensins such as NaD1 from *Nicotiana alata* and MtDef5 from *Medicago truncatula* recruit one or more phospholipids to oligomerize, induce membrane disruption, bind to intracellular targets and trigger fungal cell death (Poon *et al.*, 2014; Cools *et al.*, 2017; Islam *et al.*, 2017).

The important question related to the antifungal action of plant defensins is whether their fungicidal mechanisms are conserved in different fungi. Previously, we reported evidence that mechanisms used by MtDef4 from *M. truncatula* to inhibit the growth of *Fusarium graminearum* and *Neurospora crassa* are not the same even though these two fungi belong to the same phylum Ascomycota, subphylum Pezizomycotina and order Sordariomycetes (El-Mounadi *et al.*, 2016). This raises the possibility that the architecture and composition of the cell wall and plasma membrane may be different even in closely related fungal pathogens and markedly influence the antifungal activity of plant defensins. But it is still unknown if different developmental stages of a fungal pathogen also respond differently to a plant defensin.

To date, plant defensins that have been studied in-depth for their MOA are mostly from short-lived herbaceous annual plants (Cools *et al.*, 2017; Parisi *et al.*, 2019). To our knowledge, very few defensins from perennial woody plants have been studied for their antifungal activity and MOA. The only case is that the MOA of two members of the grapevine *(Vitis vinifera)* defensin gene family have been partially investigated (Giacomelli *et al.*, 2012; Nanni *et al.*, 2014). Olive tree *(Olea europaea)* is an evergreen woody perennial oil crop which is cultivated in Mediterranean countries for its healthy oil. The genome sequences of two cultivated and one wild ancestral olive tree (Barghini *et al.*, 2014; Cruz *et al.*, 2016; Unver *et al.*, 2017) as well as the European ash tree *(Fraxinus excelsior)* (Sollars *et al.*, 2017), all belonging to the family *Oleaceae*, have been sequenced and annotated. In this study, we have searched the genome sequences of the wild olive tree *(O. europaea* var. *sylvestris)* (Unver *et al.*, 2017), a cultivated olive tree (*O. europaea* var. *Farga*) (Cruz *et al.*, 2016) and *F. excelsior* (Sollars *et al.*, 2017) and identified several genes encoding defensins. The genomes of these trees contain a unique gene family encoding defensins not present in other eudicots.

In this study, we examined the structure-activity relationships and MOA of the antifungal defensin OefDef1.1, a member of the *O. europaea* var. *Farga* defensin family. We show that the conidia and germlings of a necrotrophic fungal pathogen *B. cinerea* respond differently to the antifungal action of this defensin. Based on our results, a model for the antifungal action of OefDef1.1 against conidia and germlings of this pathogen is proposed. A chimeric OefDef1.1 containing the γ-core motif of a plant defensin exhibits cation-resistant antifungal activity *in vitro* and confers better resistance to *B. cinerea* than the wild-type OefDef1.1. We also report a comparative analysis of the antifungal properties of this defensin against a panel of closely related ascomycete fungal pathogens in the absence and presence of elevated levels of Na^1+^.

## Results

### Identification of a novel defensin gene family in the olive tree and the European ash tree

We set out to systematically identify defensin genes in the recently published annotated genome sequences of the cultivated (*O. europaea* var. *Farga)* and wild olive tree called oleaster (*O. europaea* var. *sylvestris*) (Cruz *et al.*, 2016; Unver *et al.*, 2017), using the BLAST search as describe in MATERIALS AND METHODS. Several members of the defensin family in each olive tree were identified and phylogenetic relationships among them were determined (Fig. 1). The *O. europaea* var. *Farga* genome encodes eight defensins and the *O. europaea* var. *sylvestris* genome encodes fifteen defensins grouped into three clades. Phylogenetic analysis revealed eight homologs of MtDef4 from *Medicago truncatula* (Clade I), six homologs of MtDef5 from *M. truncatula* (Clade II) and nine defensins rich in histidine and tyrosine that are clustered in a separated Clade III on the phylogenetic tree (Fig. 1). The Clade I defensins have comparable homologs in all higher plants, whereas Clade II has comparable homologs in several other dicotyledonous plants. Of the nine defensins present in Clade III, only two (OE3B065497T1 and OE3B086095T1) are found in *O. europaea* var. *Farga*, while the remaining seven are found in *O. europaea* var. *sylvestris*. Thus, cultivated olive tree with a smaller genome also has a smaller number of the Clade III defensins. Interestingly, this novel Clade III gene family does not exist in the other plants except European ash tree *(F. excelsior)* (Sollars et al., 2017) which also belongs to the *Oleaceae* family. Four defensins sharing 83 to 88% sequence identity with one of the olive tree defensins OefDef1.1 were identified (Supplementary Fig. S1). All Clade III defensins have high predicted net charge of 7.6 to 10.3 at pH 7.0 and four predicted disulfide bonds with a conserved cysteine-stabilized αβ signature. They share more than 80% sequence identity and contain a non-canonical cationic γ-core motif which contains ten residues (GXCX_10_C), instead of the expected nine, between the two cysteine residues. Based on our analysis, we propose that Clade III defensins represent a novel *Oleaceae*-specific gene family.

**Fig. 1.**
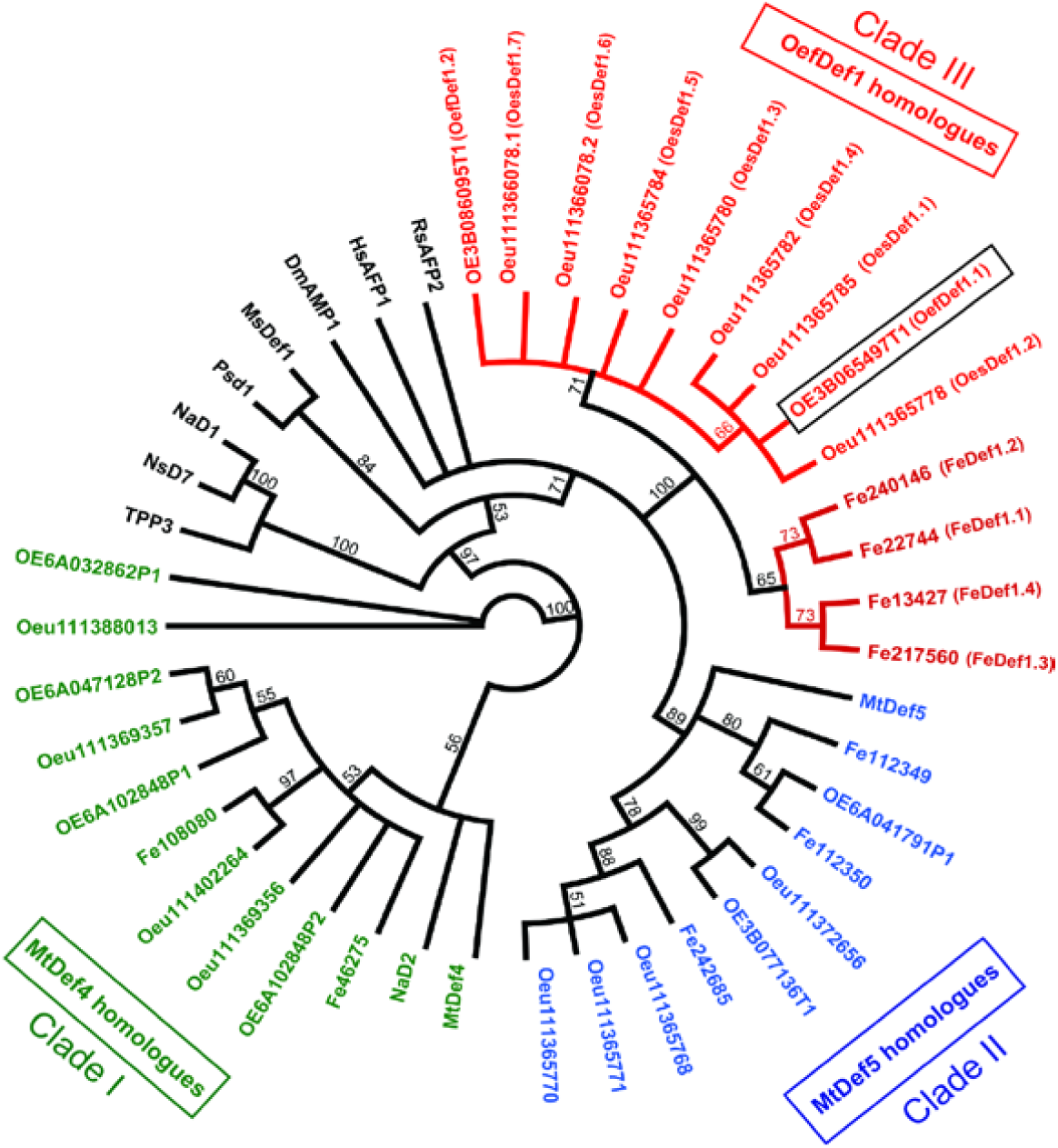
Phylogenetic tree of defensin sequences identified from olive tree. The olive tree defensin homologs were identified through BLAST on Phytozome and NCBI. Geneious software was used to generate the phylogenetic tree of defensins based on Neighbor-joining method with bootstrap (10,000 replications). Defensin sequences shown in green, blue and red color are MtDef4 homologs (Clade I), MtDef5 homologs (Clade II) and OefDef1.1 homologs (Clade III), respectively. Clade I and II defensins have corresponding homologs in other plants.

### OefDef1.1 exhibits potent antifungal activity *in vitro* and *in planta*

The amino acid sequence of OefDef1.1 (OE3B065497T1) harboring a net charge of 9.3 is shown in Supplementary Fig. S1. *In vitro* antifungal activity of this peptide against *B. cinerea* and three *Fusarium* spp. was determined. It inhibited the growth of these fungi with the IC_50_ values of 1.6±0.6 µM for *F. graminearum*, 1.1±0.2 µM for *F. virguliforme*, 0.4±0.1 µM for *F. oxysporum* and 0.7±0.3 µM for *B. cinerea* (Table 1). Growth inhibition of *B. cinerea* was also observed under the microscope which showed that treatment of *B. cinerea* conidia with 1.5 µM OefDef1.1 completely inhibited its germination (Fig. 2A). In addition, resazurin cell viability assay was used to determine the concentration at which each peptide caused 100% cell death (minimum inhibitory concentration, MIC). Thus, cell death was observed at a concentration of 1.5 µM OefDef1.1 in *B. cinerea* (Fig. 2B). These results indicated that OefDef1.1 has potent broad-spectrum antifungal activity.

**Table 1.**
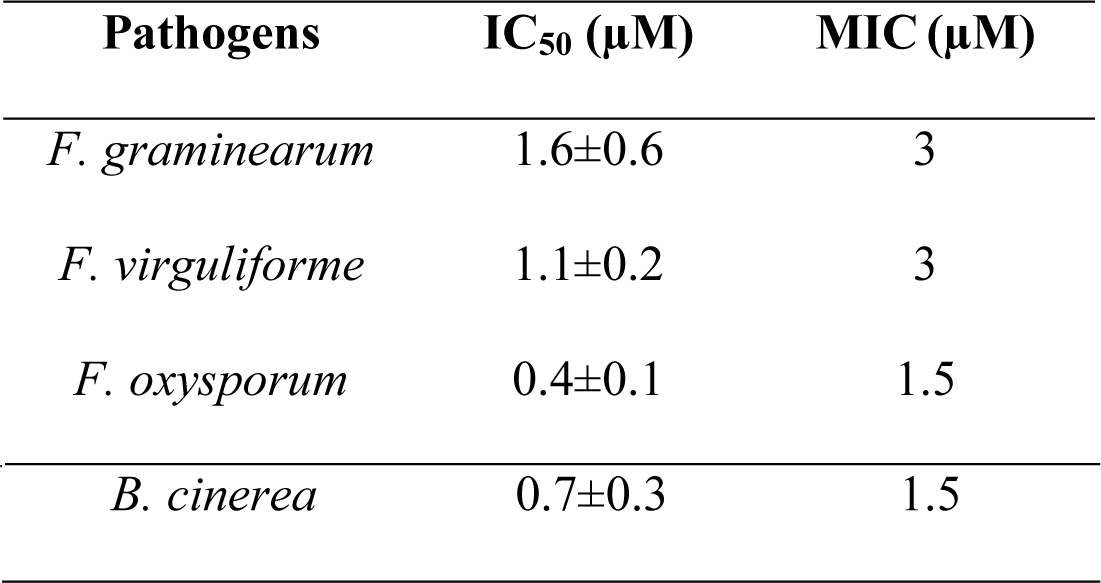
IC_50_ and MIC values of Oefdef1.1 against fungal pathogens. IC_50_ values were calculated from the inhibition curve for each fungal pathogen. Values are mean±SD; n=3. MIC values were determined by the resazurin cell viability assay.

**Fig. 2.**
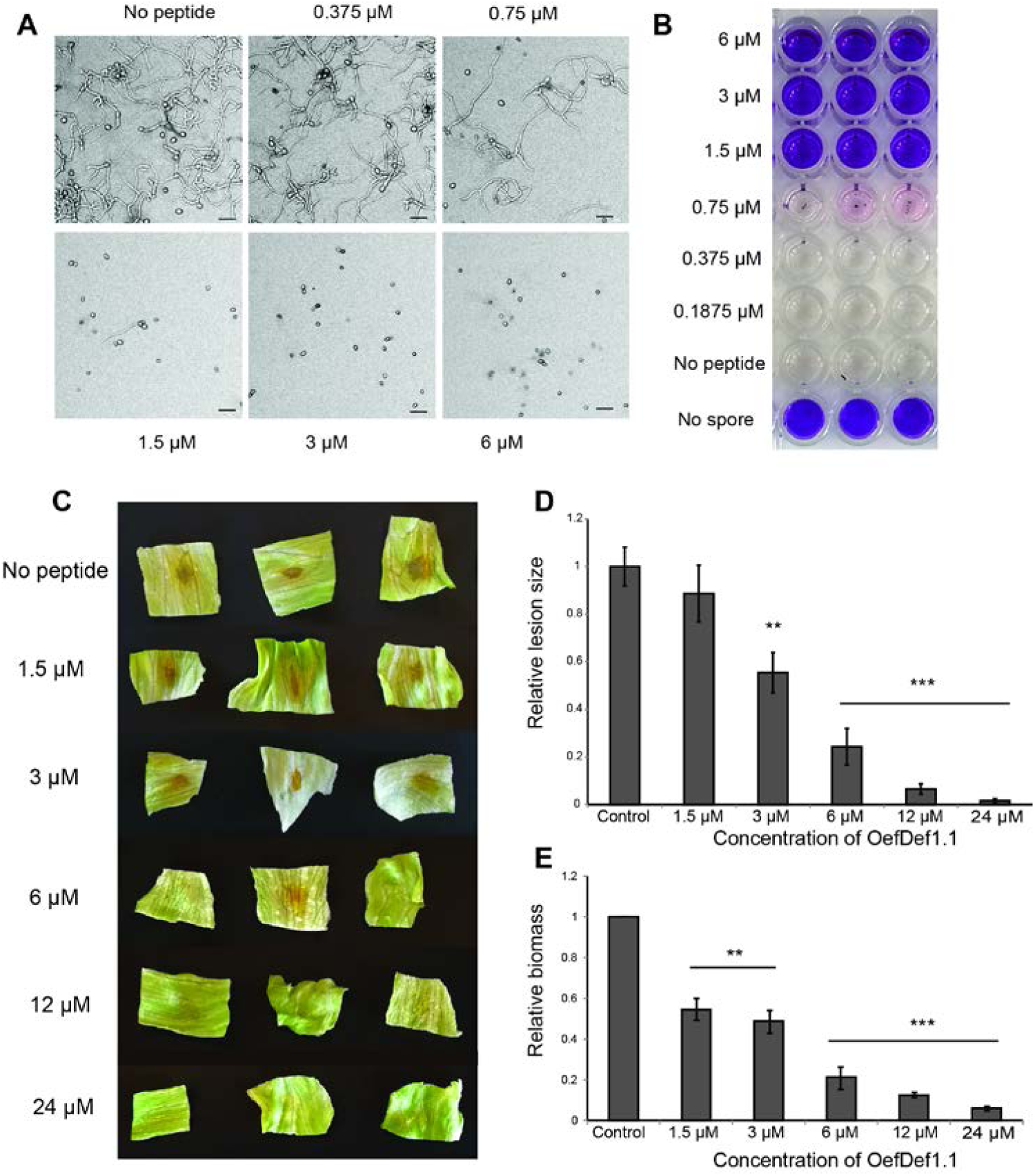
OefDef1.1 shows broad-spectrum antifungal activity *in vitro* and reduces grey mold disease symptoms *in planta.* A. Micrographs displaying growth inhibition of *B. cinerea* at different concentrations of OefDef1.1 are shown. Complete inhibition of spore germination was observed at a concentration of 1.5 µM. Bar = 40 µm. Images were taken at 60 hpi. B. Fungal cell death in *B. cinerea* was assessed using the resazurin cell viability assay. The wells with dark blue color indicated absence of live cells. C. Lesions caused by *B. cinerea* infection of the detached lettuce leaves were progressively reduced in size with increasing concentrations of OefDef1.1. Leaves with no peptide applied were used as controls. Images were taken at 48 hpi. D. Relative lesion sizes at different concentrations of OefDef1.1 were measured by ImageJ software. E. Relative fungal biomass indicated by the relative fungal DNA content was determined by qPCR. Leaves without peptide treatment were used as controls. The statistical analysis was performed using *t* test to determine whether the observed differences were statistically significant (**, P< 0.01; ***, P< 0.001). Error bars denote standard deviations.

Since OefDef1.1 inhibits the growth of *B. cinerea in vitro*, we evaluated its ability to reduce the symptoms of the grey mold disease *in planta*. Different concentrations of the peptide were applied on the surface of the detached lettuce leaves followed immediately with the inoculum of fresh *B. cinerea* conidia. After 48 h, the leaves treated with no peptide control showed large lesions; however, higher concentrations of peptides produced significantly smaller lesions of fungal infection (Fig. 2C and D) and the 6 µM OefDef1.1 almost completely prevented development of disease lesions. We also determined the relative DNA content of *B. cinerea* by quantitative PCR. This analysis confirmed a significant decrease in the biomass of *B. cinerea* at a peptide concentration of 1.5 µM, although a more pronounced decrease in the biomass was observed at a peptide concentration over 6 µM (Fig. 2E). Thus, surface application of OefDef1.1 efficiently reduces symptoms of *B. cinerea* infection in lettuce.

### OefDef1.1 rapidly permeabilizes the plasma membrane of the conidia and germlings of *B. cinerea*

Since *B. cinerea* is an economically important pathogen responsible for causing extensive pre- and post-harvest decay in fruits, vegetables and flowers (Dean *et al.*, 2012), we have chosen it as a model fungus to elucidate the MOA of OefDef1.1.

Killing of fungal cells by antifungal plant defensins often correlates with the permeabilization of the plasma membrane. We used SG uptake assay to determine the effect of OefDef1.1 on the plasma membrane integrity in real-time in the cells of *B. cinerea*. Fresh conidia and germlings were incubated with 3 µM OefDef1.1 and SG uptake was monitored continuously by time-lapse confocal microscopy. In fresh conidia, the SG uptake was apparent within 4 min and increased with time (Fig. 3). After 20 min, the nuclei were completely stained with SG, indicating rapid permeabilization of the conidial cells by OefDef1.1. OefDef1.1 also permeabilized the plasma membrane of germlings rapidly. SG signal was observed in germlings as early as 1 min following the peptide challenge (Fig. 3).

**Fig. 3.**
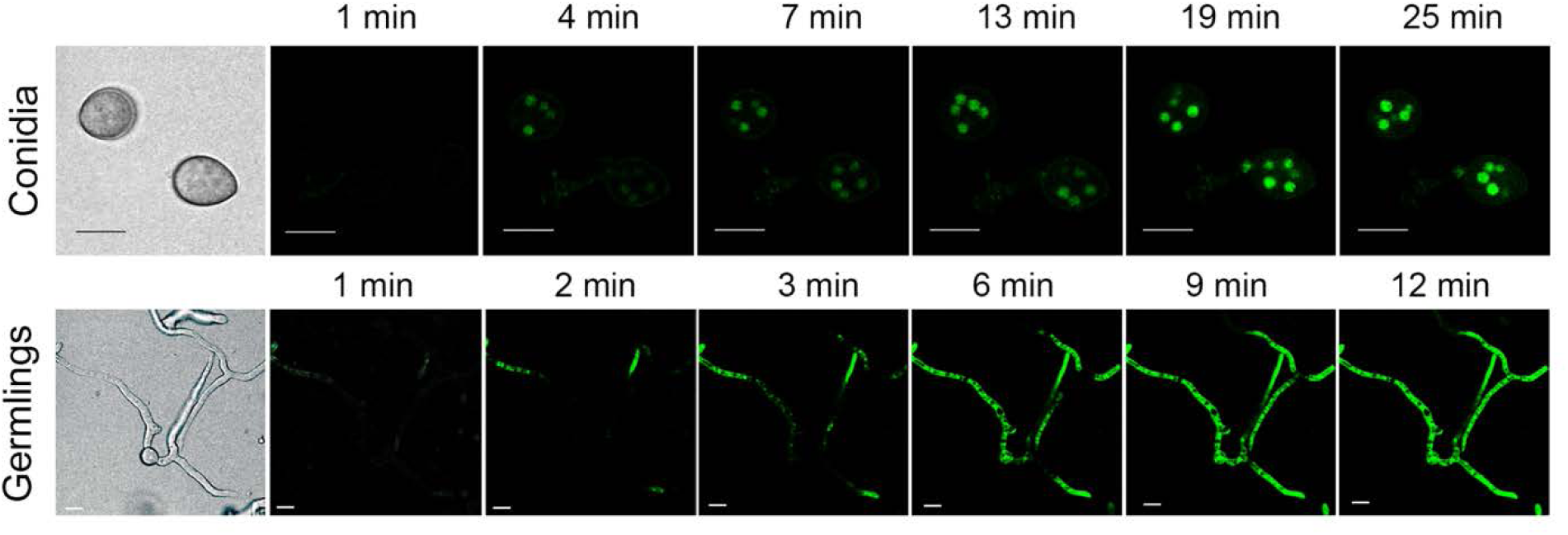
OefDef1.1 induces membrane permeabilization both in conidia and germlings of *B. cinerea*. *B. cinerea* conidia and germlings were incubated with 3 µM OefDef1.1 and 1 µM SYTOX Green (SG). Images under confocal microscopy were taken every 3 min for conidia and every 1 min for germlings after OefDef1.1 challenge. Bars= 10 μm.

### Subcellular localization of OefDef1.1 and ROS production are different in the conidia and germlings of *B. cinerea*

We sought to determine if antifungal activity of OefDef1.1 is initiated by binding to the cell wall followed by translocation to cytoplasm in the conidia and germlings of *B. cinerea*. OefDef1.1 labeled with the fluorophore DyLight 550 was added to freshly harvested conidia. As shown in Fig. 4A, OefDef1.1 bound to the cell wall within 10 min of challenge, and it remained on the cell wall even after 130 min of challenge. If labeled peptide was added to conidia along with the membrane selective dye FM4-64, the DyLight 550 signal did not co-localize with the FM4-64 signal (Supplementary Fig. S2). No DyLight 550 labeled OefDef1.1 signal was detected in the cytoplasm of the conidia, indicating that in freshly harvested conidia OefDef1.1 acts via the extracellular side.

**Fig. 4.**
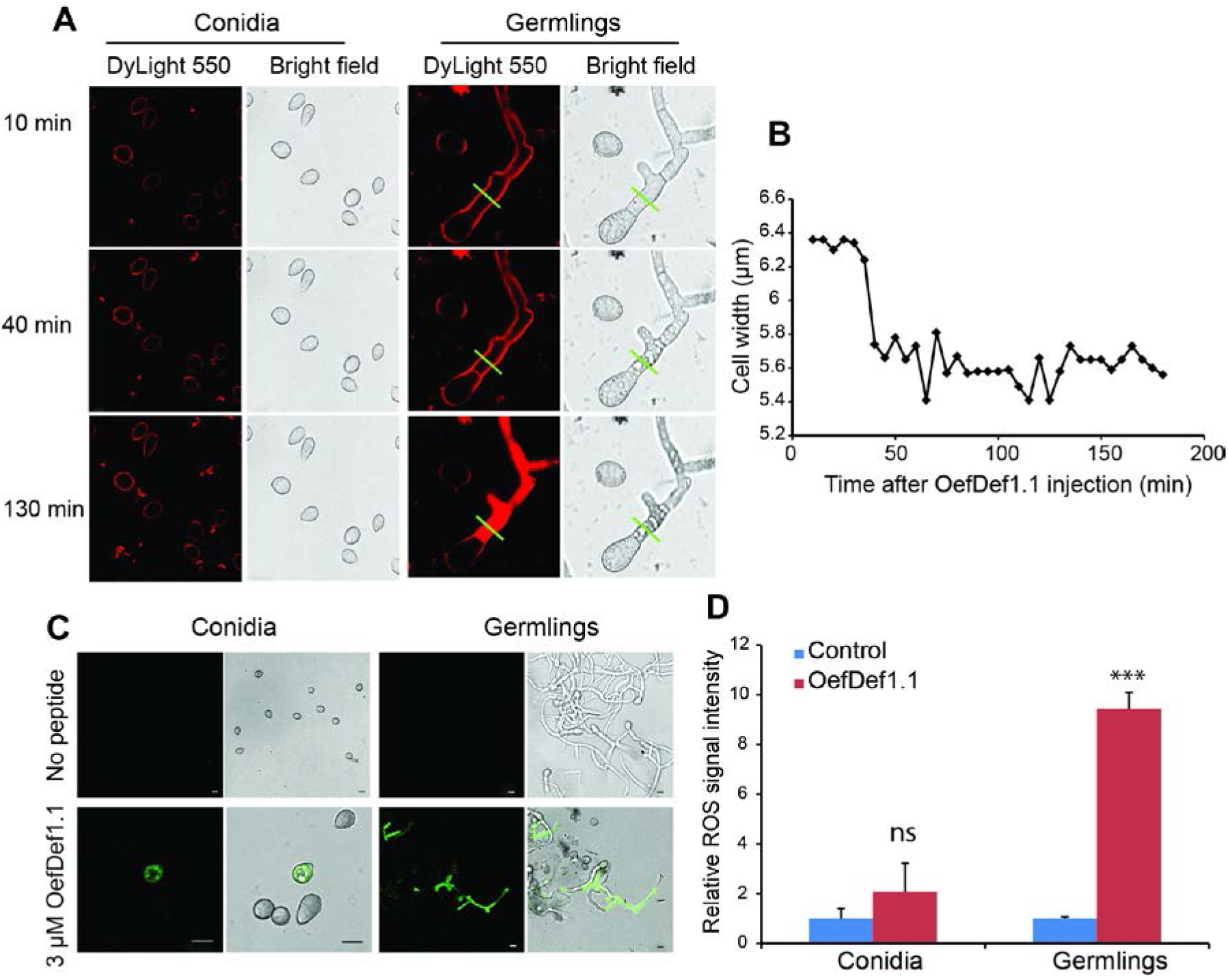
Uptaking OefDef1.1 and ROS production are different in conidia and germlings of *B. cinerea*. A. The intracellular localization of 12 μM DyLight 550 labeled OefDef1.1 in *B. cinerea* conidia and germlings. The confocal microscope images were captured every 5 min after OefDef1.1 challenge. Images taken at 10 min, 40 min and 130 min are shown. B. Cell width was determined by quantifying the inner distance between cell walls in brightfield images from germling cell shown in A (the measuring location is labeled by green line). C. Confocal microscope images showing ROS production (green fluorescence) in fresh conidia and germlings after treatment with 3 μM OefDef1.1 for 90 min. Bars = 10 μm. D. The relative ROS signal intensity in fresh conidia and germlings. Samples without OefDef1.1 treatment were set as control. The statistical analysis was performed by *t* test to determine whether the observed differences were statistically significant (P<0.001) as shown by asterisks. ns= not significant (P> 0.05). Error bars denote standard deviation values.

DyLight 550 labeled OefDef1.1 bound to the cell wall of *B. cinerea* germlings 10 min after challenge as similar as conidia (Fig. 4A). However, after 40 min of challenge, the color of cytoplasm became dark and germlings shrunk suddenly, and the distance between cell walls decreased from 6.2 nm to 5.6 nm accompanied by shrinkage in the cell volume (Fig. 4A and B). After 40 min, DyLight 550 labeled OefDef1.1 began to enter into the cytoplasm of germling cells and, at 130 min, filled the cells uniformly (Fig. 4A).

The induction of oxidative stress by antifungal peptides contributes to cell death in fungi. We determined if OefDef1.1 induces ROS production in the conidia and germlings of *B. cinerea.* In conidia and germlings, ROS signal was observed following the OefDef1.1 treatment for 90 min. However, a significant difference in the ROS levels was observed in the two cell types (Fig. 4C and D). The ROS signal did not accumulate significantly in fresh conidia upon OefDef1.1 challenge but increased nine-fold in germlings (Fig. 4D). These results indicate that germlings are more sensitive to the oxidative stress induced by OefDef1.1 than conidia and that ROS likely play a role in killing of germlings but not conidia.

### Homology-based three-dimensional structure of OefDef1.1 and analysis of its γ-core substitution variants

Using the solution structure of *Artemisia vulgaris* Artv1 defensin (PDB: 2KPY), a homology model of the three-dimensional structure of OefDef1.1 has been generated (Fig. 5A). As expected, the predicted structure of OefDef1.1 contains the structural characteristics found in other plant defensins for which structural information exists.

**Fig. 5.**
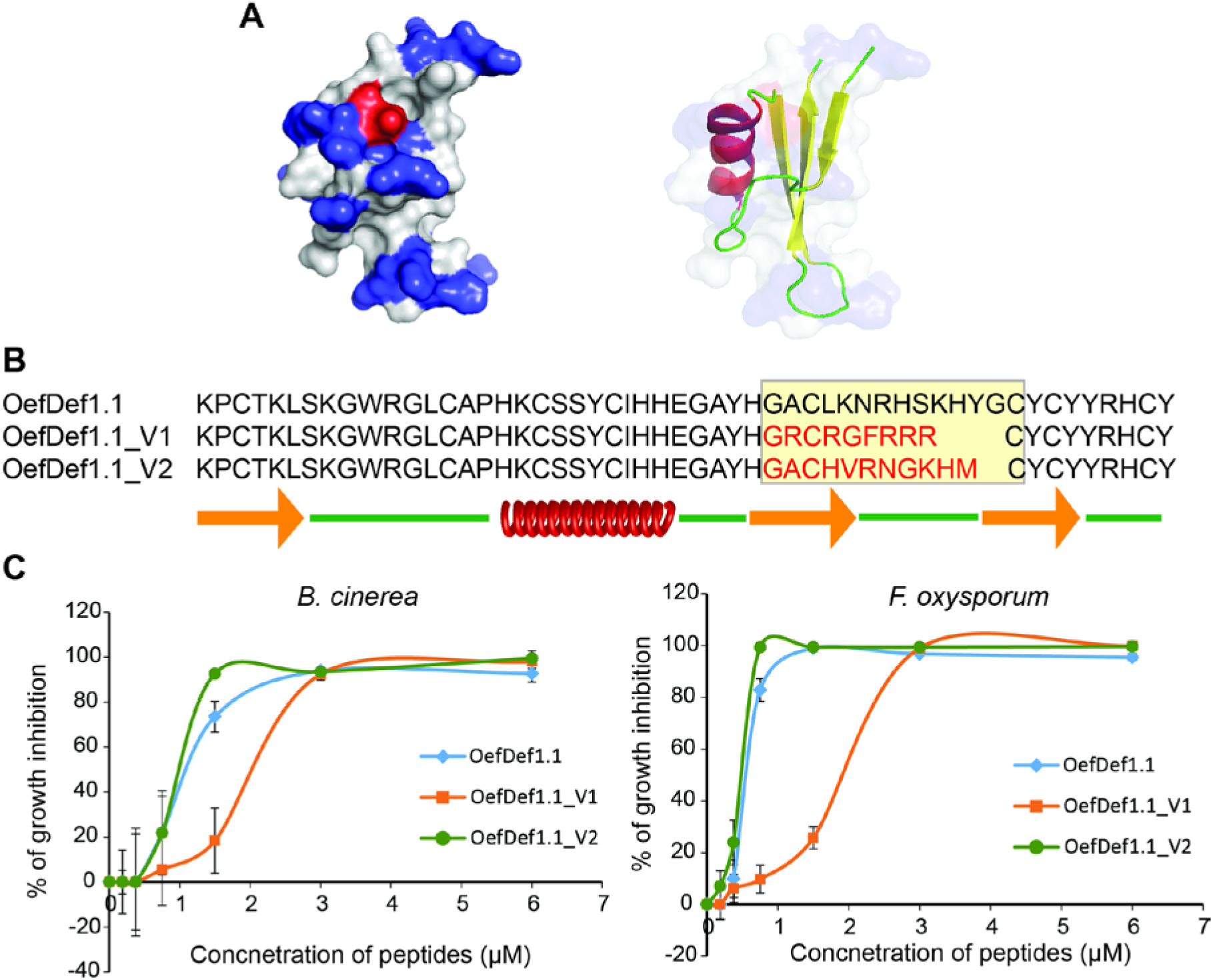
Homology-based three-dimensional structure and antifungal activity of its γ-core motif variants. A. 3D structure and the solvent-exposed surface of the wild-type OefDef1.1 were constructed by homology-based modeling. The surface representation of OefDef1.1 (left) displays the cationic residues in blue, anionic residues in red, and uncharged or hydrophobic residues in white. The cartoon representation (right) displays α-helix in red, β-sheets in yellow and loops in green. The *Artemisia vulgaris* Artv1 (PDB number: 2KPY) was used as the template. B. The amino acid sequences of the wild-type OefDef1.1 and its variants. The γ-core motifs of OefDef1.1 and its two variants are shown in the yellow rectangle, and γ-core motif sequences selected for substitution are shown in red. α-helix is shown as a red spiral, three β-sheets are shown as orange arrows, and the loops are shown as green lines, respectively. C. Antifungal activity of the wild-type OefDef1.1 and its variants against *B. cinerea* and *F. oxysporum*. Error bars denote standard deviation values.

As reported, the highly conserved γ-core motif plays an important role in the antifungal activity, phospholipid binding and oligomer formation of a plant defensin. To determine as to what extent the γ-core motif contributes to the antifungal activity of OefDef1.1, we generated two variants, OefDef1.1_V1 and OefDef1.1_V2, in which this motif was replaced by the corresponding γ-core motif of MtDef4 or that of DmAMP1 from *Dahlia merckii*, respectively (Fig. 5B and supplementary Table S1). The γ-core motif of MtDef4 (GRCRGFRRRC) is very different from that of OefDef1.1 and is more cationic. DmAMP1 sequence is 41% similar to the OefDef1.1 sequence, but its γ-core motif is different in sequence from that of OefDef1.1.

Antifungal activity of the variants against *B. cinerea* and *F. oxysporum* was determined (Fig. 5C). OefDef1.1_V1, which contains the γ-core motif of MtDef4, lost about 50% activity compared with the wild-type OefDef1.1. It inhibited the growth of these fungi with an IC_50_ value of 2-3 µM and MIC value of 4-6 µM. In contrast, OefDef1.1_V2 containing the γ-core motif of DmAMP1 showed higher antifungal activity than the wild-type OefDef1.1 with an MIC value of 1.5 µM.

### OefDef1.1_V1 is more effective than the wild-type OefDef1.1 in reducing symptoms of *B. cinerea* infection *in planta*

Earlier, we showed that the wild-type OefDef1.1, when applied topically on the detached leaves of iceberg lettuce, significantly inhibited grey mold disease caused by *B. cinerea* (Fig. 2C-E). We decided to compare *in planta* antifungal activity of OefDef1.1_V1 and OefDef1.1_V2. Another variant OefDef1.1_V5 with *in vitro* antifungal activity similar to that of OefDef1.1 (unpublished data) was also included in this analysis for comparison. Because of their large size and susceptibility to *B. cinerea*, detached leaves *N. benthamiana* were used in this assay. The wild-type OefDef1.1 and variants at different concentrations were applied topically to the surface of the leaf and infected immediately with the conidia of *B. cinerea*. Both lesion size and photosynthetic efficiency were measured. To avoid leaf-to-leaf variation, all four peptides along with no peptide control were spotted on the surface of the same leaf. Based on the Fv/Fm value, the condition of each infected leaf was rated as healthy (Fv/Fm > 0.7, shown in green), slightly damaged (Fv/Fm 0.5-0.7, shown in yellow) or severely damaged (Fv/Fm < 0.5, shown in red) (Supplementary Fig. S3). When compared with no peptide control, each peptide reduced the size of lesions caused by infection of the pathogen (Fig.6A and B). As expected, the percentage of damaged leaves including slightly damaged and severely damaged parts decreased with increasing concentration of each peptide (Fig. 6C). Among these four tested peptides, OefDef1.1_V2 showed similar efficacy in reducing disease lesions as the wild-type OefDef1.1, although the Fv/Fm data suggested that OefDef1.1_V2 was more potent at lower concentrations (3 µM and 6 µM). Surprisingly, OefDef1.1_V1 with much reduced antifungal activity *in vitro* was most effective in inhibiting the grey mold disease symptoms. At 3 µM OefDef1.1_V1, 80% reduction in lesion size was observed and, at a concentration of 6 µM, disease symptoms were completely attenuated (Fig. 6A). Similar results were also observed in lettuce (Fig. 6D and E). Thus, OefDef1.1_V1 with higher net charge is most effective in controlling grey mold disease than peptides with lower net charge.

**Fig. 6.**
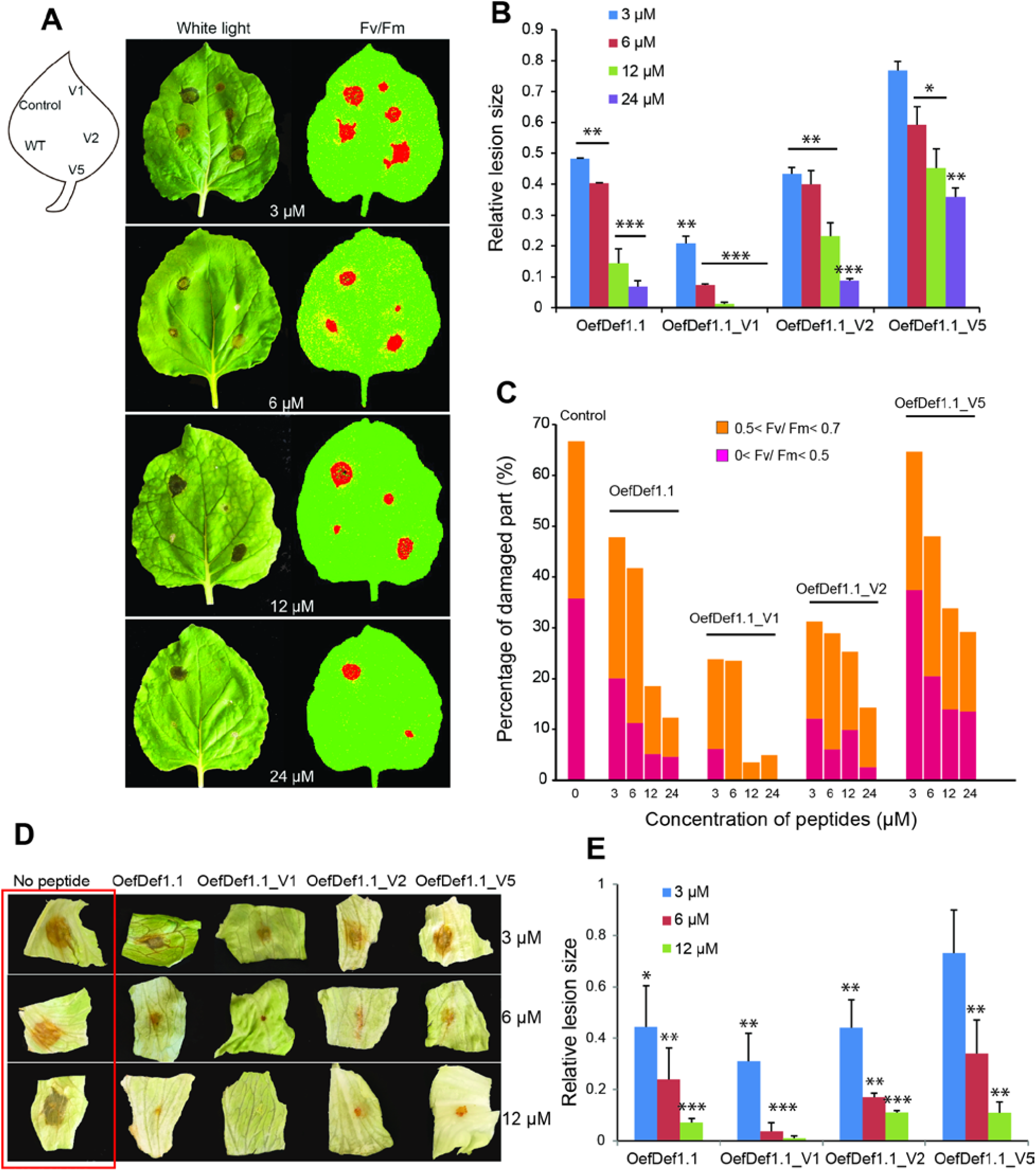
The inhibition of grey mold disease symptoms on the *Nicotiana benthamiana* and lettuce leaves by topical application of OefDef1.1 and variants. A. The lesions formed by *B. cinerea* were imaged under white light and high resolution images taken by CropReporter by determining the value of potential photosynthetic efficiency (Fv/Fm). The value of Fv/Fm was shown as different colors (red, Fv/Fm <0.5; yellow, 0.5< Fv/Fm < 0.7; green, Fv/Fm > 0.7). Different concentrations (0, 3 µM, 6 µM, 12 µM and 24 µM) of OefDef1.1, OefDef1.1_V1, OefDef1.1_V2 and OefDef1.1_V5 were applied onto the leaves first and fresh conidia of *B. cinerea* were applied at the same spot. Images were captured after 48 hpi. B. The relative lesion size was measured from the white light images using the ImageJ software. The lesion formed without peptide was used as a control. C. The percentage of damaged part of each 35*35 pixels frame containing the whole lesion of each treatment. Fv/Fm values for the red and orange columns are indicated. The lesions caused without peptide treatment were set as control (100%). D. Lesions formed by *B. cinerea* on lettuce leaves applied with 3 µM, 6 µM and 12 µM of OefDef1.1, OefDef1.1_V1, OefDef1.1_V2 and OefDef1.1_V5. The control samples are highlighted by a red rectangle. E. The relative lesion sizes on lettuce leaves as determined by ImageJ software. The control sample with no peptide applied was set as 1. Statistical analysis was performed by *t* test to determine whether the observed differences were statistically significant (*, P<0.05; **, P<0.01; ***, P< 0.001). Error bars denote standard deviation value. Three independent replications were performed for each experiment.

### Antifungal activity of OefDef1.1_V1 is more tolerant to mono- and divalent cations than that of OefDef1.1

Antifungal activity of plant defensins is significantly reduced in buffers containing mono- and divalent cations potentially limiting their efficacy as antifungal agents *in vivo*. It has been hypothesized that the presence of cations significantly weakens the electrostatic interactions between a positively charged defensin and negatively charged fungal membranes (Melo *et al.*, 2009). We therefore evaluated the antifungal activity of the wild-type OefDef1.1 and its more positively charged OefDef1.1_V1 in the presence of SFM buffer (a low salt medium) supplemented with 100 mM NaCl, 100 mM KCl or 1 mM CaCl_2_. As expected, antifungal activity the wild-type OefDef1.1 is significantly reduced in presence of all three cations. Even at a 4 x MIC concentration of 6 µM OefDef1.1, significant conidial germination was observed in presence of the cations (Fig. 7A). OefDef1.1_V1 behaved differently from that of OefDef1.1. In contrast to the wild-type OefDef1.1, antifungal activity of this variant was enhanced in presence of 100 mM NaCl or 100 mM KCl, and nearly no decrease in its antifungal activity occurred in presence of 1 mM CaCl_2_. These results were confirmed by the resazurin cell viability assay (Supplementary Fig. S4A). The cationic media used in this study had no deleterious effects on the growth of *B. cinerea* (Supplementary Fig. S4B). These results indicate that increased cationicity positively correlates with the higher antifungal activity of the peptide in presence of cations.

**Fig. 7.**
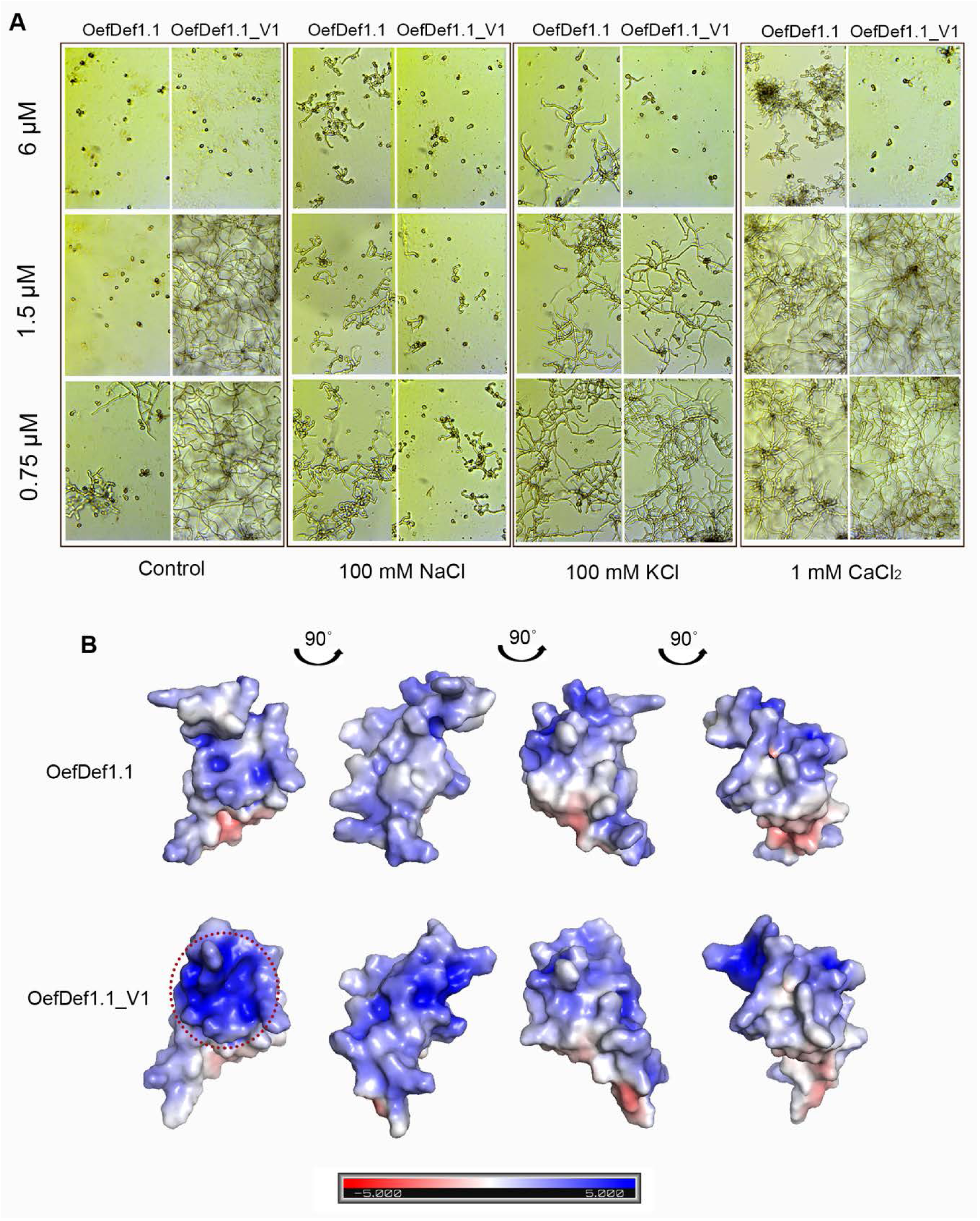
Antifungal activity of OefDef1.1 and OefDef1.1_V1 against *B. cinerea* in the presence of mono- and bivalent cations and surface charge of these peptides. A. The growth of *B. cinerea* treated with OefDef1.1 and OefDef1.1_V1 in media containing cations. Images were captured by microscopy at 60 hpi. B. Qualitative electrostatic surface representation of OefDef1.1 and OefDef1.1_V1 (blue=cationic residues, red=anionic residues and white=uncharged or hydrophobic residues). The cationic pocket is highlighted by a red dotted circle. The surface charges are based on the dielectric constant at ±5 (bar).

A comparison of the solvent accessible surface potential plots of the 3D structures of OefDef1.1 and OefDef1.1_V1 indicated the presence of a highly charged pocket similar to that recently reported for a maize defensin ZmD32 (Kerenga *et al.*, 2019) in the variant, but not in the wild-type OefDef1.1 (Fig.7B). The formation of a cationic pocket on the protein surface might be responsible for the enhanced antifungal activity of OefDef1.1_V1 in presence of Na^1+^.

### Antifungal properties of OefDef1.1 vary significantly among closely related ascomycete fungal pathogens in presence of cations

OefDef1.1 loses its antifungal activity against *B. cinerea* in presence of cations, but whether this cations-sensitive property is conserved among the other fungal pathogens is not clear. We therefore decided to examine the antifungal mechanisms of OefDef1.1 in presence of cations against four ascomycete fungal pathogens, *F. graminearum, F. virguliforme, F. oxysporum* (subphylum Pezizomycotina and class Sordariomycetes) and *B. cinerea* (subphylum Pezizomycotina and class Leotiomycetes).

We examined the ability of OefDef1.1 to permeabilize the plasma membrane and translocate into the interior of the conidia and germlings of each pathogen in absence and presence of 100 mM NaCl. In addition, we determined the antifungal activity of OefDef1.1 against each pathogen as determined by 100% inhibition of conidial germination. In medium without elevated Na^1+^, OefDef1.1 permeabilized the plasma membrane of conidia in all pathogens. As expected, additional Na^1+^significantly reduced the ability of OefDef1.1 to permeabilize the plasma membrane of conidia in each pathogen, but quantitative differences were observed among pathogens (Table 2, Supplementary Fig. S5). In low salt medium, both *B. cinerea* and *F. virguliforme*, more than 90% of conidia were permeabilized by OefDef1.1. However, in presence of additional Na^1+^, only 3.9% of *B. cinerea* conidia were permeabilized while 35.8% of *F. virguliforme* conidia could still be permeabilized by this defensin.

**Table 2.**
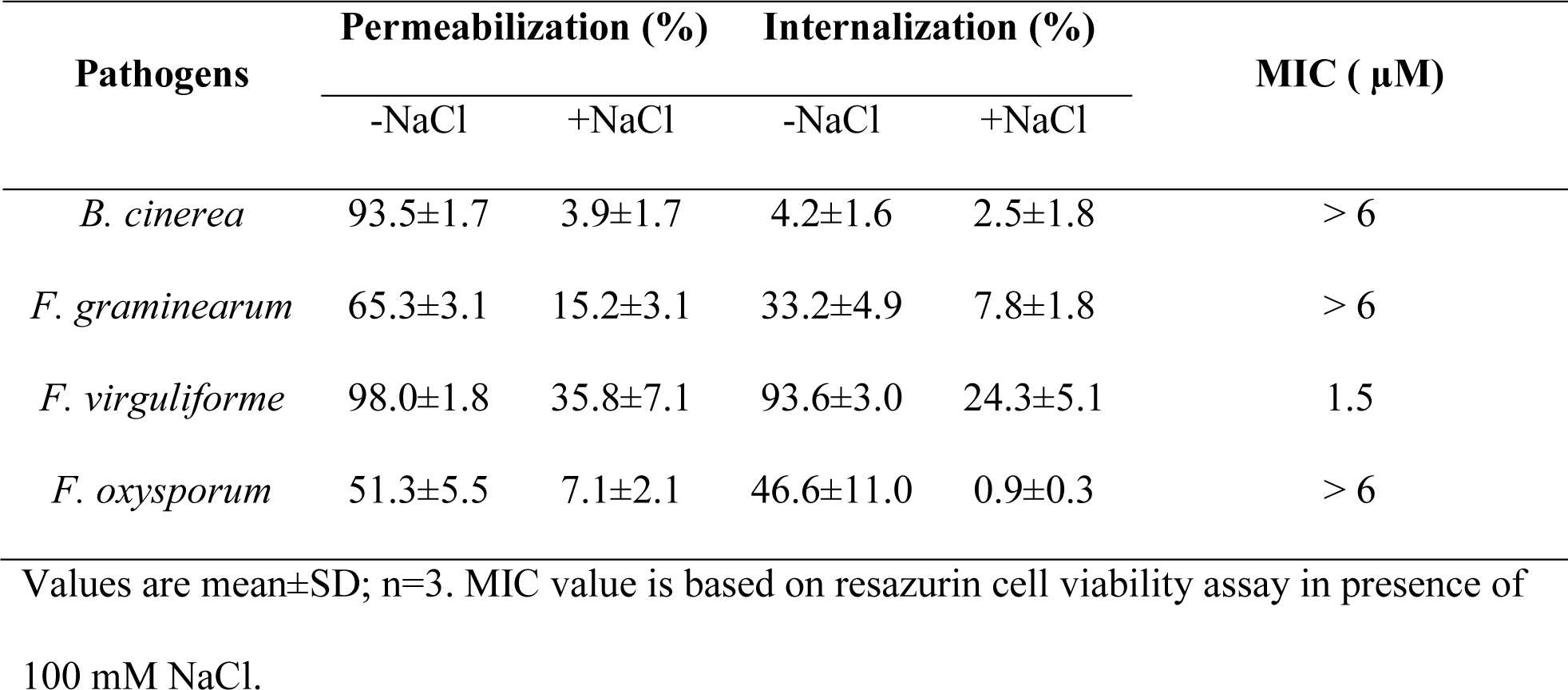
Percentage of plasma membrane permeabilization and defensin internalization in conidia from different fungal pathogens.

| Pathogens | Permeabilization (%) |  | Internalization (%) |  | MIC ( μM) |
| --- | --- | --- | --- | --- | --- |
|  | -NaCl | +NaCl | -NaCl | +NaCl |  |
| *B. cinerea* | 93.5±1.7 | 3.9±1.7 | 4.2±1.6 | 2.5±1.8 | > 6 |
| *F. graminearum* | 65.3±3.1 | 15.2±3.1 | 33.2±4.9 | 7.8±1.8 | > 6 |
| *F. virguliforme* | 98.0±1.8 | 35.8±7.1 | 93.6±3.0 | 24.3±5.1 | 1.5 |
| *F. oxysporum* | 51.3±5.5 | 7.1±2.1 | 46.6±11.0 | 0.9±0.3 | > 6 |

Values are mean±SD; n=3. MIC value is based on resazurin cell viability assay in presence of 100 mM NaCl.

Significant quantitative differences were also noted in the internalization of OefDef1.1 into conidia of these pathogens in the absence of elevated Na^1+^ (Table 2, Supplementary Fig. S5). In medium without additional Na^1+^, the conidia of three *Fusarium* spp., but not of *B. conidia*, internalized OefDef1.1. In presence of elevated Na^1+^, uptake of OefDef1.1 was inhibited in all four pathogens, although 24.3% of *F. virguliforme* conidia still retained the ability to internalize the peptide. However, in *F. oxysporum* conidia, internalization of the peptide was completely blocked in presence of high Na^1+^. Thus, the conidia of the two *Fusarium* spp. responded differently to the antifungal action of OefDef1.1 in presence of a high concentration of the monovalent cation. Consistent with these results, antifungal activity of OefDef1.1 was retained against *F. virguliforme* in presence of Na^1+^, but not against the other three fungi (Supplementary Fig. S6, Table 2).

The ability of OefDef1.1 to permeabilize the plasma membrane and enter into cytoplasm of germilings was almost completely blocked in presence of high Na^1+^ in *B. cinerea, F. graminearum* and *F. oxysporum*. However, in *F. virguliforme* germlings, the uptake of SG and DyLight 550 labeled peptide was still clearly visible (Fig. 8). Based on these results, we conclude that the antifungal properties of OefDef1.1 can vary even in closely related fungal pathogens.

**Fig. 8.**
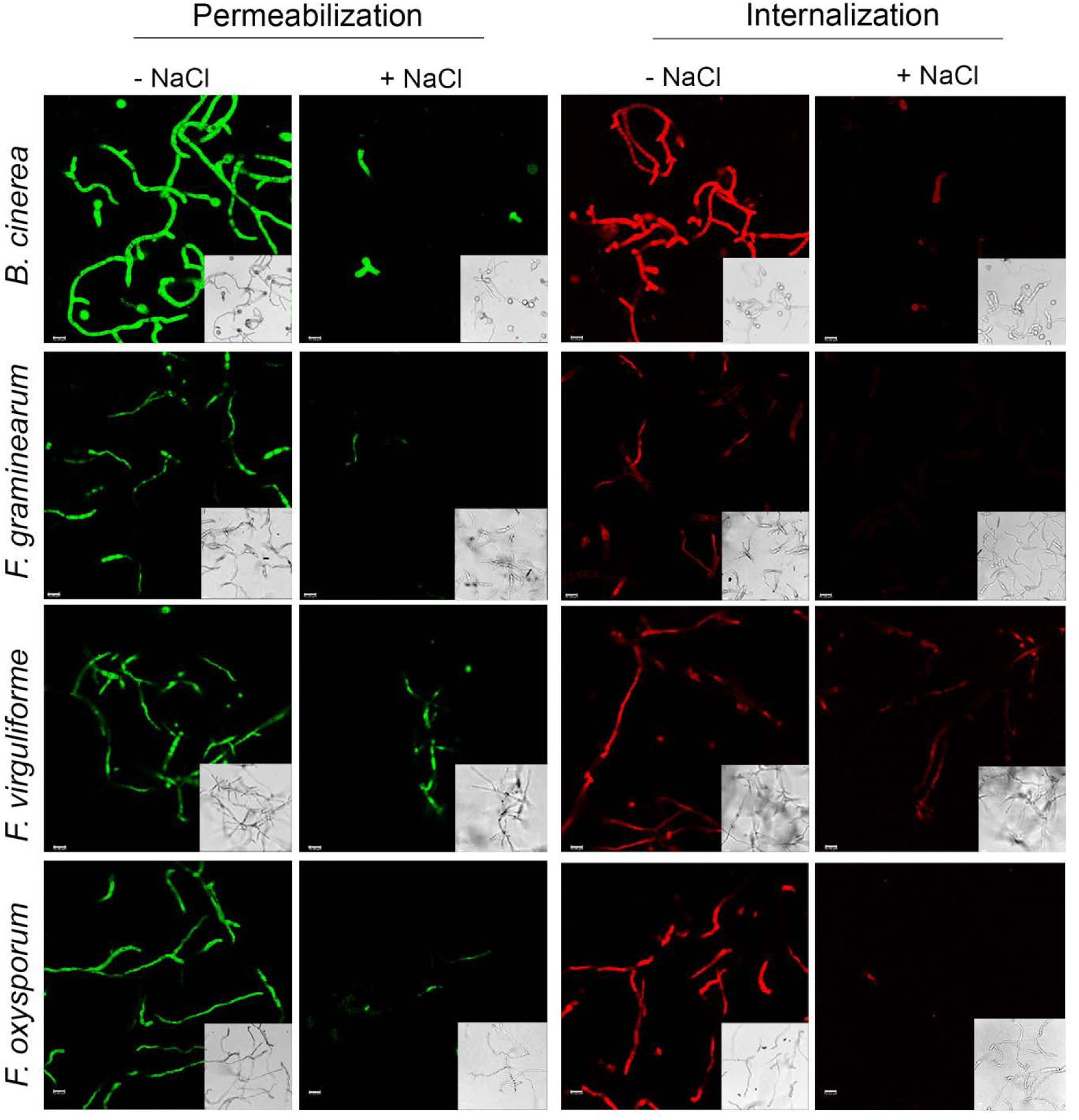
Plasma membrane permeabilization of fungal cells and internalization of OefDef1.1 into fungal cells in different pathogens incubated with or without 100 mM NaCl. Germlings of each fungal pathogen were treated with 2x MIC values of peptide. Images were captured after 3 h of defensin challenge. SG was used at 1 µM. Bars = 10 μm.

## Discussion

In this study, we have identified a novel gene family encoding highly cationic histidine- and tyrosine rich defensins in the olive tree, an evergreen perennial woody plant. This family of defensins appears to be unique to the family *Oleaceae*. Since the annotation of the olive tree and the European ash tree draft genomes is still far from complete, the exact number of these genes in the genome of each tree remains to be determined with certainty. Nine defensins in the olive tree share more than 90% sequence identity and thus appear to have evolved as a result of recent gene duplication. The uniqueness of these defensins also suggests that they have likely evolved to perform specific biological functions in the *Oleaceae* family. They are among the most positively charged peptides in the plant kingdom and contain a high percentage of hydrophobic residues. However, they each contain a cationic γ-core motif different in sequence and length from those in other well characterized defensins.

In the present study, we have found that OefDef1.1 exhibits potent broad-spectrum antifungal activity against four closely related ascomycete fungal pathogens and it significantly reduces the symptoms of grey mold disease caused by *B. cinerea* infection *in planta*. The ability of this defensin to inhibit fungal pathogen growth *in vitro* and *in planta* leads us to predict that this defensin plays a role in host defense against fungal pathogens in olive tree.

Antifungal plant defensins vary in their MOA. However, they all share a common characteristic of interacting with the fungal cell wall and permeabilizing the plasma membrane of fungal pathogens (Cools *et al.*, 2017; Parisi *et al.*, 2019). Very little is known at present regarding the cell wall components interacting with plant defensins. In our mechanistic studies employing *B. cinerea* as our model fungus, we have determined that OefDef1.1 quickly binds to the cell walls of conidia and germlings. However, in the presence of 100 mM NaCl, it still binds to the cell wall of conidia, but very weakly to that of germlings indicating its cell-type specific interaction with the cell wall.

Fungal cell wall is a dynamic organelle and its structure and composition display significant variations between different cell types and growth conditions within the same species (Geoghegan *et al.*, 2017). Thus, the relative concentrations of β1,3-glucan, β1,6-glucan, chitin, mannan and glycoproteins, the major components of the cell wall, might vary between cell types. Significant changes in the major polysaccharides of the cell wall occur when a *B. cinerea* conidium germinates and differentiates into a germling (Ruiz-Herrera, 1992; Cantu *et al.*, 2009). Significant changes in the carbohydrate composition of the cell wall at different stages (spores, mycelium and zygotes) of development have also been reported in *Penicillium roqueforti* and *Absidia coerulea* (Andriyanova *et al.*, 2011). Fungal cell wall also comprises a complex matrix of interconnected polysaccharides and proteins (Latge, 2007). Thus, differences in the cell wall proteomes of the conidia and germlings could account for cell-type specific differences observed in the cell-wall binding of OefDef1.1 in presence of a monovalent cation Na^1+^.

Our data have identified notable differences among three *Fusarium* spp. and *B. cinerea* with regard to OefDef1.1’s ability to permeabilize the plasma membrane and gain entry into conidial and germling cells and kill fungal cells in absence and presence of elevated Na^1+^. It is particularly interesting that, in presence of high Na^1+^, OefDef1.1 permeabilizes the plasma membrane and enters the conidial and germling cells of *F. virguliforme*, but not of closely related *F. oxysporum*. These observations are noteworthy especially since the two *Fusarium* spp. belong to the same *Nectriaceae* family and exhibit hemi-biotrophic lifestyles. Although molecular targets of OefDef1.1 in the cell wall of fungal pathogens used in this study are not yet known, significant differences must exist in the composition and architecture of the cell wall of closely related fungal species of the phylum Ascomycota to account for varied responses of the fungal species to OefDef1.1 (Geoghegan *et al.*, 2017).

OefDef1.1 rapidly permeabilizes the plasma membrane of *B. cinerea*, suggesting that it is a critical event in the fungicidal action of this defensin. Membrane permeabilization has been shown to be an important early step in the MOA of plant defensins such as NaD1 (Poon *et al.*, 2014) and MtDef5 (Islam *et al.*, 2017). These defensins engage phoshoinositides or related phospholipids to oligomerize and induce membrane permeabilization (Parisi *et al.*, 2019).

OefDef1.1 shows very weak or no binding to phospholipids in the protein-lipid overlay and liposome binding assays (Fig. S7). Therefore, the mechanism by which membrane permeabilization is induced by OefDef1.1 remains to be defined. Induction of ROS is hypothesized to be another important event in the antifungal action of plant defensins including NaD1 (van der Weerden *et al.*, 2008), RsAFP2 (Aerts *et al.*, 2007) and MtDef5 (Islam *et al.*, 2017). OefDef1.1 challenge also induces ROS production in germlings, suggesting their involvement in cell death. However, the ROS signal intensity induced by OefDef1.1 in freshly harvested conidia only accounts for 20% of the ROS signal in germlings, suggesting that oxidative stress might be a significant factor in the killing of germlings, but not conidia.

Several plant defensins have now been shown to be translocated inside the fungal cells (Cools *et al.*, 2017; Parisi *et al.*, 2018). It has been proposed that these defensins bind to intracellular targets as part of their MOA. Internalization of OefDef1.1 appears to be cell type-dependent and varies among fungal species. In *B. cinerea*, OefDef1.1 remains bound to the cell wall of conidia and almost no uptake (less than 5%) is observed even after 3 h of challenge. However, it is internalized rapidly in the germlings of this pathogen. In contrast, this defensin is internalized in both cell types of the fungal pathogens, *F. virguliforme* and *F. oxysporum*. To what extent internalization of this peptide contributes to the killing of fungal cells in these pathogens remains to be determined. It is certainly not required for killing of conidial cells in *B. cinerea*.

The structure of OefDef1.1 is similar to those of other plant defensins. However, its γ-core motif (GACLKNRHSKHYGC) is different from those of other well characterized plant defensins in length and sequence. The γ-core motif contains major determinants for phospholipid binding, oligomer formation and antifungal activity of several plant defensins (Sagaram *et al.*, 2012; de Coninck *et al.*, 2013; Cools *et al.*, 2017; Parisi *et al.*, 2019). The analysis of the *in vitro* and *in planta* antifungal activity of two chimeric OefDef1.1 variants each containing a heterologous γ-core motif yielded unexpected results. OefDef1.1_V1 containing the γ-core motif sequence (GRCRGFRRRC) of MtDef4 was less active against *B. cinerea* than the wild-type OefDef1.1 *in vitro*, but it provided greater control of this pathogen *in planta*. In contrast, OefDef1.1_V2 containing the γ-core motif sequence (CHVRNGKHMC) of DmAMP1 was more active than the wild-type OefDef1.1 *in vitro*, but provided less control of this pathogen *in planta*. Thus, under the assay conditions used, *in vitro* antifungal activity of a plant defensin does not necessarily correlate with *in planta* antifungal activity. One likely explanation for this apparent discrepancy is that OefDef1.1_V1 exhibits greater antifungal activity in presence of cations than either OefDef1.1 or OefDef1.1_V2. The ability to retain antifungal activity in presence of cations may be important for a plant defensin to confer resistance to a fungal pathogen *in planta*. Recently, highly cationic maize defensin ZmD32 with broad-spectrum antifungal activity in media containing elevated levels of Na^1+^ has been characterized (Kerenga *et al.*, 2019). As in OefDef1.1_V1, ZmD32 also contains a positively charged pocket in its 3D structure. The RGFRRR motif present in the γ-core motif of each defensin is responsible the formation of this cationic pocket and cation tolerant antifungal activity.

Based on the MOA studies reported here, we propose a multistep model for the antifungal action of OefDef1.1 against *B. cinerea* (Fig. 9). Although no difference was observed in the MIC value of this defensin between conidia and germlings (data not shown), their interactions with this peptide were significantly different. In both conidia and germlings, the first two steps are initial binding to the cell wall followed by rapid disruption of the plasma membrane. Na^1+^ blocks cell wall binding of the peptide in germlings. While in conidia Na^1+^ does not block binding of the peptide to the cell wall, but it does block membrane permeabilization. After 90 min of peptide challenge, significantly more ROS is detected in germlings than in conidia. Within 3 h of peptide challenge, massive internalization of the peptide and significant cell shrinkage is observed in germlings, but not in conidia. Better understanding of the relative contribution of each step to the antifungal action of this defensin will require further studies. However, rapid membrane permeabilization stands out as a major contributing factor.

**Fig. 9.**
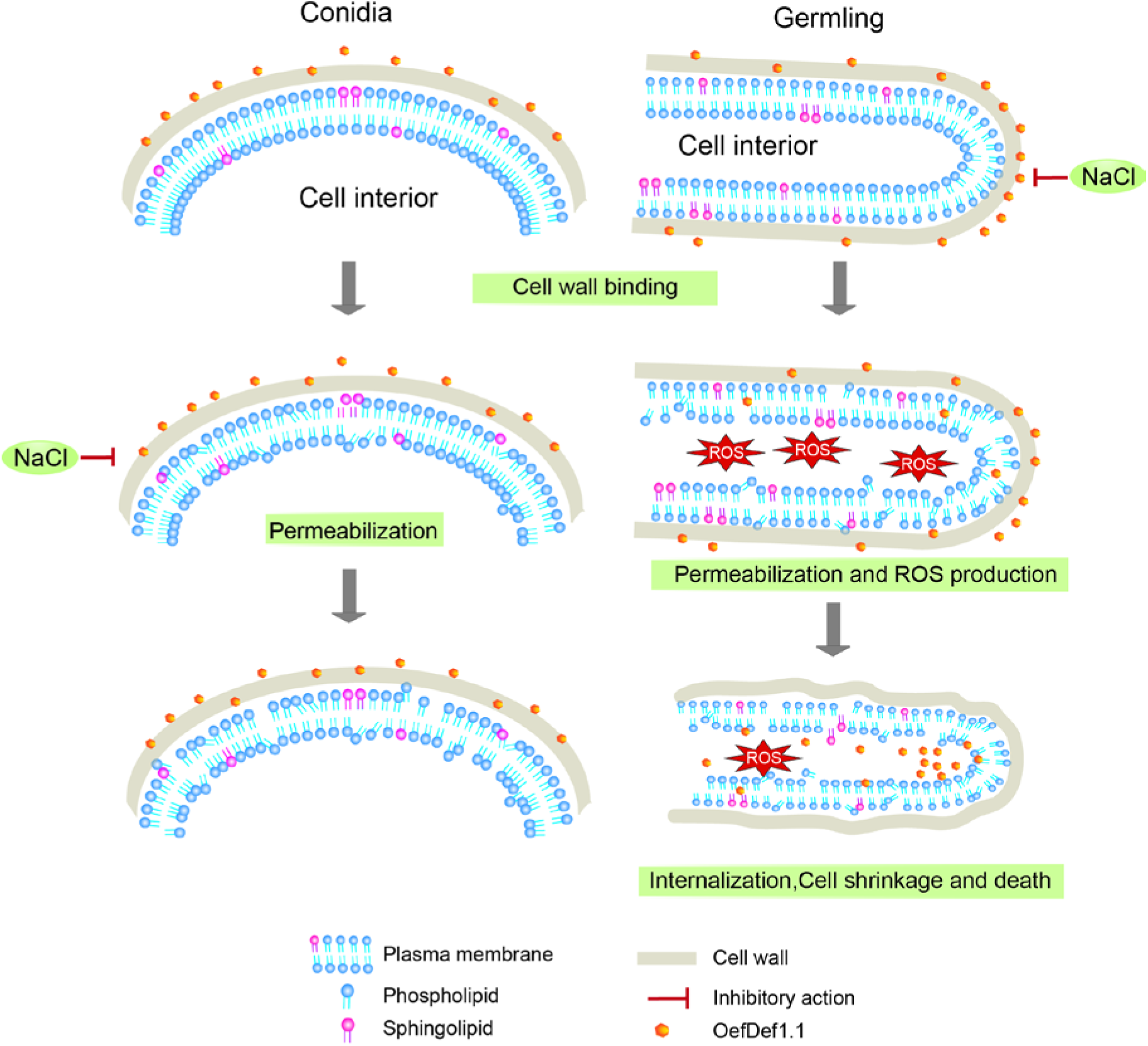
Proposed models showing differences in the fungicidal action of OefDef1.1 against the conidia and germlings of *B. cinerea*. OefDef1.1 binds to the cell wall and permeabilizes the plasma membrane of conidia and germlings. However, NaCl blocks cell wall binding of the peptide in germlings, but not in conidia. It also blocks membrane permeabilization in conidia. After 90 min of peptide challenge, significantly more ROS is detected in germlings than in conidia. Within 3 h of peptide challenge, massive internalization of the peptide and significant cell shrinkage is observed in germlings, but not in conidia.

## MATERIALS AND METHODS

### Fungal cultures and growth medium

The fungal strains *Fusarium graminearum* PH-1, *F. virguliforme, F. oxysporum* f. sp. *Cubense* and *Botrytis cinerea* strain T-4 were cultured in media shown in Supplementary Table S2.

### Defensin gene identification and phylogenetic analysis

Defensin genes were identified through BLASTP using the default parameters on Phytozome, a plant genomics resource portal, and NCBI databases using as queries peptide sequences of MtDef4 and MtDef5 (Sagaram *et al.*, 2011; Islam *et al.*, 2017). All other defensin sequences were obtained from the available published literature. The mature peptide sequences were aligned using multiple sequence alignment tool MUSCLE, and the phylogenetic tree was inferred by the Neighbor-Joining method using the Geneious software with bootstrap (10,000 replications).

### Recombinant expression and purification of OefDef1.1 and its variants in *Pichia pastoris*

The codon-optimized OefDef1.1, OefDef1.1_V1 and OefDef1.1_V2 genes were synthesized by GenScript (Piscataway, NJ). The genes for OefDef1.1 and its variants were each expressed in *P. pastoris* and the peptide was purified from the culture medium using the Fast Protein Liquid Chromatography System (GE Healthcare) and Reverse Phase-High Performance Liquid Chromatography (RP-HPLC) and subsequently lyophilized as described (Islam *et al.*, 2017). Each peptide was re-suspended in nuclease-free water and its concentration was determined by either NanoDrop spectrophotometry or the BCA assay. Purity and size of each peptide were determined by electrophoresis on a 4-20% Mini-Protean TGX gels (Bio-Rad). The correct mass of each peptide was confirmed by mass spectrometry prior to its use in experiments described below.

### *In vitro* Antifungal activity assays

Antifungal activity assays were conducted as described previously with minor modifications (Spelbrink *et al.*, 2004). The quantitative fungal growth inhibition by OefDef1.1 and its variants was estimated by measuring the absorbance at 595 nm using a Tecan Infinite M200 Pro (Tecan Systems Inc., San Jose, CA) microplate reader at 48 h. The fungal/oomycete cell viability/cell killing was determined by the resazurin cell viability assay (Chadha & Kale, 2015). After incubation of the pathogen/peptide mixture for 48 h, 10 µl of 0.1% resazurin solution was added to each well and re-incubated overnight. A change in the color of the resazurin dye from blue to pink or colorless indicated the presence of live fungal cells.

Effect of cations on the antifungal activity of OefDef1.1 and OefDef1_V1 against *B. cinerea* conidia was tested as described above in presence of 100 mM NaCl or 100 mM KCl or 1 mM CaCl_2_. The plates were incubated at room temperature for 60 h and images were taken by microscopy (Leica DMI 6000B).

### Antifungal activity of OefDef1.1 and its variants *in planta*

Detached leaf infection assays were performed as described with minor modifications (Wang *et al.*, 2016). Briefly, leaves of iceberg lettuce and *Nicotiana benthamiana* were placed in Petri dishes. A 10 μl aliquot containing different concentrations of each defensin was placed onto the leaf samples and inoculation with *B. cinerea* was initiated at the same spot by applying 10 μl of spore suspension at a final concentration of 10^5^ spores ml^-1^. Petri dishes were kept in Ziploc WeatherShield plastic boxes containing wet paper towel at room temperature for 48 h. Lesions were photographed and the relative lesion size was determined using ImageJ software. The biomass of fungal pathogen was determined by quantitative PCR with primers shown in Table S2. The high resolution images of the *B. cinerea* infection of the *N. benthamiana* leaves were obtained using CropReporter (PhenoVation, Netherlands). The chlorophyll fluorescence and Fv/Fm (variable fluorescence over saturation level of fluorescence) images of the infected leaves were captured using the fluorescence imaging technology (Gorbe *et al.*, 2015).

### Plasma membrane permeabilization

Membrane permeabilization of fungal cells was analyzed using confocal microscopy by visualizing the influx of the fluorescent dye SYTOX Green (SG) (Thermo Fisher Scientific). Fresh conidia or germlings mixed with 2 x MIC (3 µM for *B. cinerea* and *F. oxysporum* and 6 µM *for F. graminearum* and *F. virguliforme)* OefDef1.1 and 1 µM SG were deposited onto glass-bottom petri dishes and imaged by confocal microscopy at an excitation wavelength of 488 nm and an emission wavelength ranging from 520 nm to 600 nm at specific time intervals. Control plates with SG but without OefDef1.1 were used as negative controls. A Leica SP8-X confocal microscope was used for all confocal imaging.

### Uptake of OefDef1.1 by fungal cells

OefDef1.1 was labeled with DyLight 550 amine reactive dye following the protocol provided by the manufacturer (Thermo Fisher Scientific). Time-lapse confocal laser scanning microscopy was performed to monitor uptake and subcellular localization of the fluorescently labeled peptide as described previously (Islam *et al.*, 2017). Since labeled OefDef1.1 lost 50% of its antifungal activity, it was used at a final concentration of 6 µM for *B. cinerea* and *F. oxysporum* or 12 µM for *F. graminearum* and *F. virguliforme*. For co-localization assay, DyLight 550-labeled OefDef1.1 was added to *B. cinerea* conidia along with 5 µM membrane selective dye FM4-64. The excitation and emission wavelength for DyLight 550 were 562 nm and 580-680 nm and for FM4-64 690 and 800 nm, respectively.

### Intracellular ROS detection

Intracellular ROS were detected in fresh conidia and 16 h-old germlings after exposure to 3 µM OefDef1.1 for 90 min. After treatment, 2′,7′-dichlorodihydrofluorescein diacetate (DCFH-DA, Sigma-Aldrich) was added at a final concentration of 10 μM and live cell imaging was performed using the Leica SP8-X microscope as described previously (Islam *et al.*, 2017).

## Acknowledgments

We thank Drs. Shin-Cheng Tzeng and Bradley Evans of the Proteomics and Mass Spec Facility for conducting mass spec analysis of OefDef1.1 and its variants. We thank Dr. Howard Berg for his expert guidance and help with confocal microscopy. We are deeply grateful to Intrexon Company for supporting this research.

