## Supplementary material for "Antifungal potency and modes of action of a novel olive tree defensin against closely related ascomycete fungal pathogens": FigS1-S7, Table S1-S3

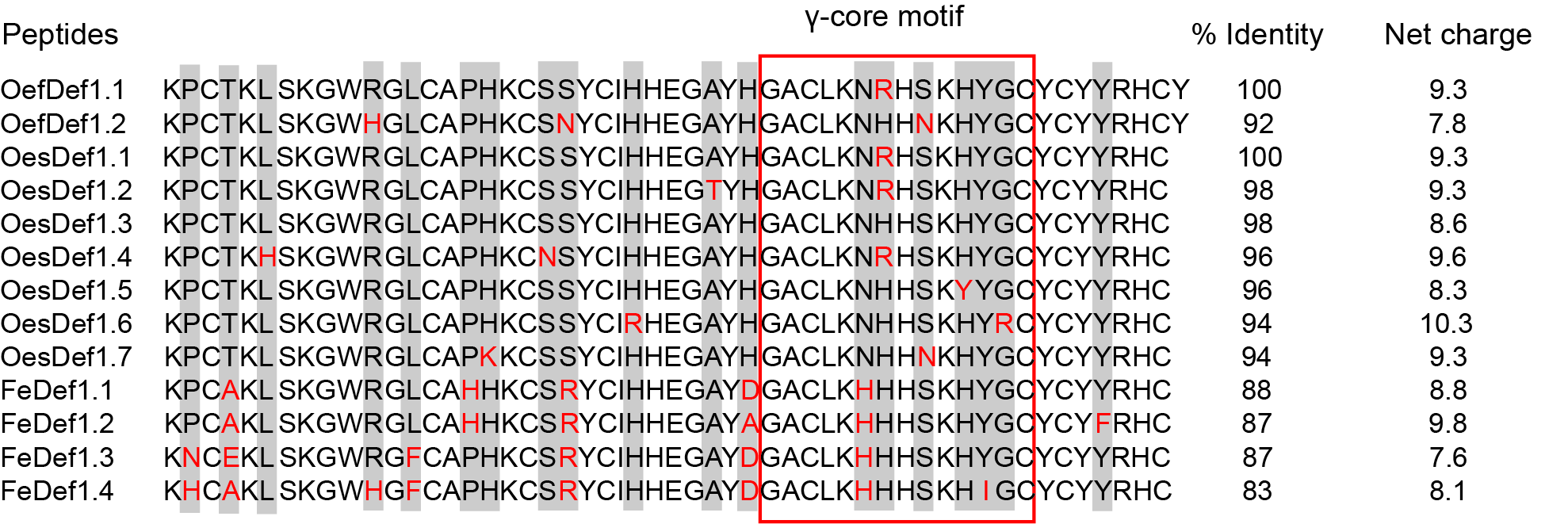

**Fig. S1 Amino acid sequence alignment of OefDef1.1 homologs.** Protein sequence alignment of OeDef1.1 and its homologs from the cultivated olive tree *O. europaea* var. *Farga* (OefDef1.2), the wild olive tree *O. europaea* var. *sylvestris* (OesDef1.1 to OesDef1.7) and ash tree *F. excelsior* (FeDef1.1-1.4). The positions of amino acid residues not conserved in all defensin sequences are shaded gray. Residues in red are the amino acid substitutions. The γ-core motif in each defensin is shown within a red rectangle. Net charge of each defensin is calculated at pH 7.0.

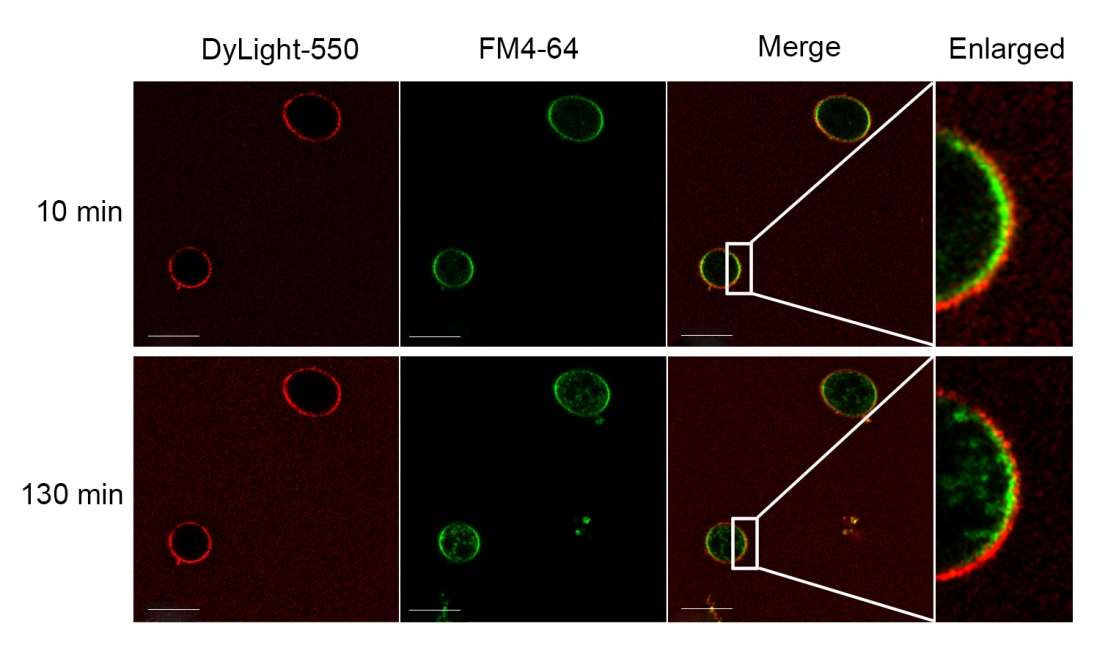

**Fig. S2** DyLight 550 labeled OefDef1.1 does not co-localize with membrane dye FM4-64 in fresh *B. cinerea* conidia. FM4-64 dye binds to cell surface within 10 min and enters into cytoplasm within 130 min, whereas DyLight 550 labeled OefDef1.1 remains localized to the cell surface and the two signals do not overlap. Bars=10 µm.

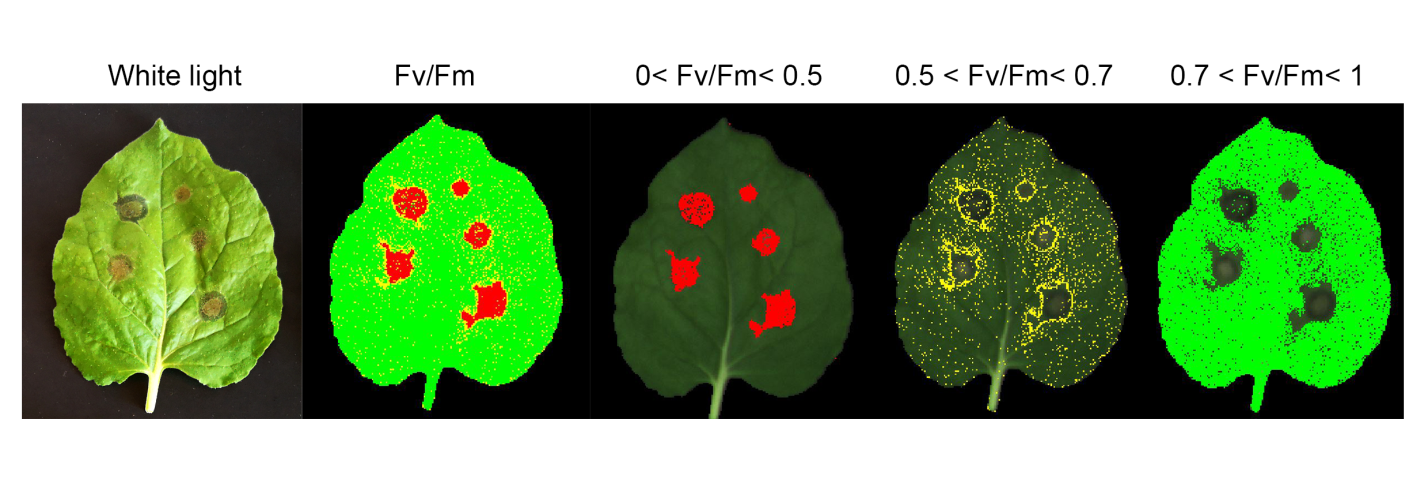
**Fig. S3** The infection lesions of *Nicotiana benthamiana* leaves by *B. cinerea* were classified as severe infection (0< Fv/Fm<0.5, red dots), mild infection (0.5< Fv/Fm<0.7, yellow dots) and no infection (Fv/Fm>0.7, green dots).

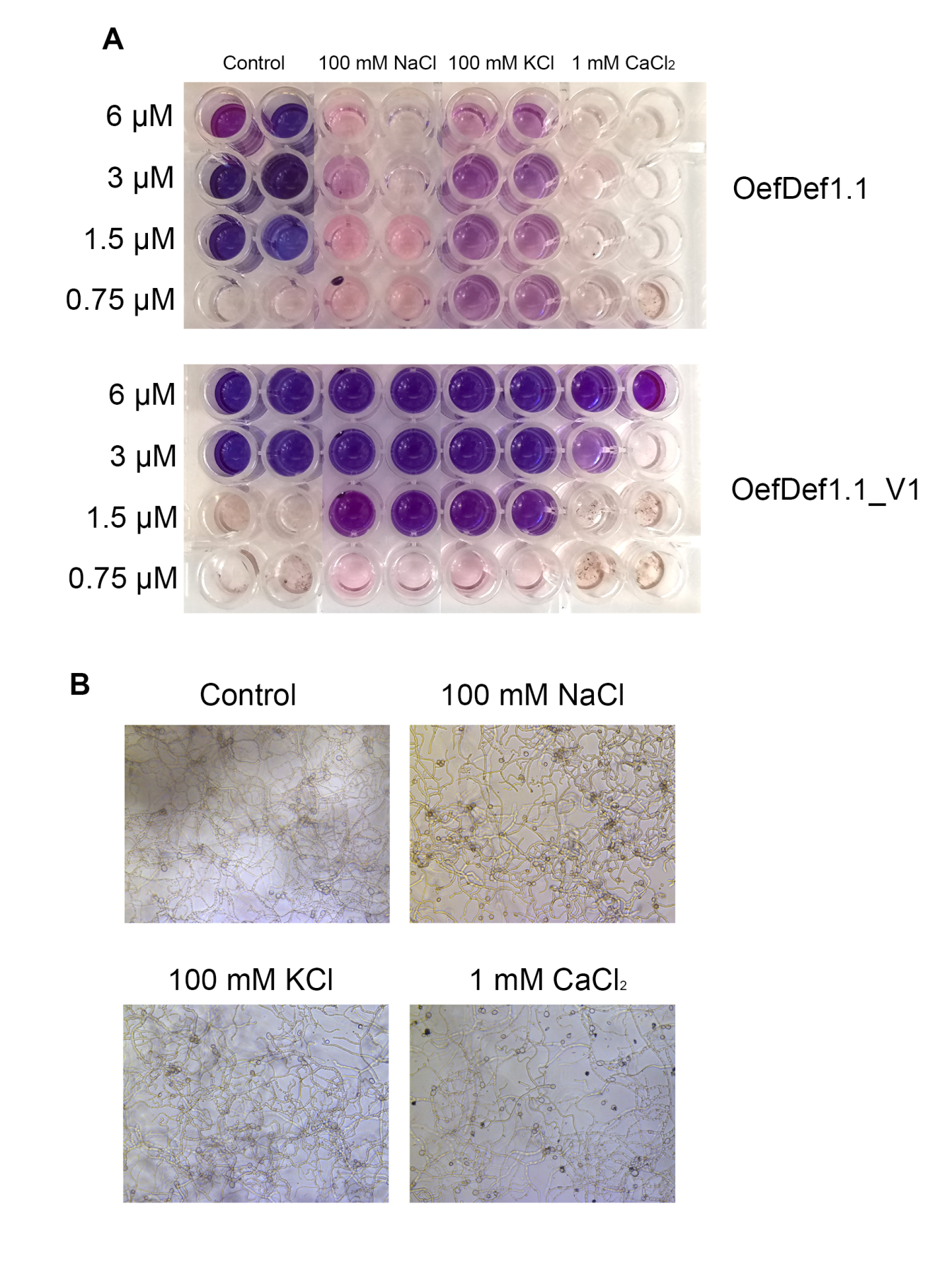

**Fig. S4** The effect of cations on the antifungal activity of OefDef1.1 and OefDef1.1_V1. AThe antifungal activity of OefDef1.1 and OefDef1.1_V1 was tested in the presence of different cations using the resazurin cell viability assay. Dark blue color indicated all cells have been killed and pink color or no color indicated cells were still alive. B. Cations used alone without peptide showed no influence on mycelial growth in this assay.

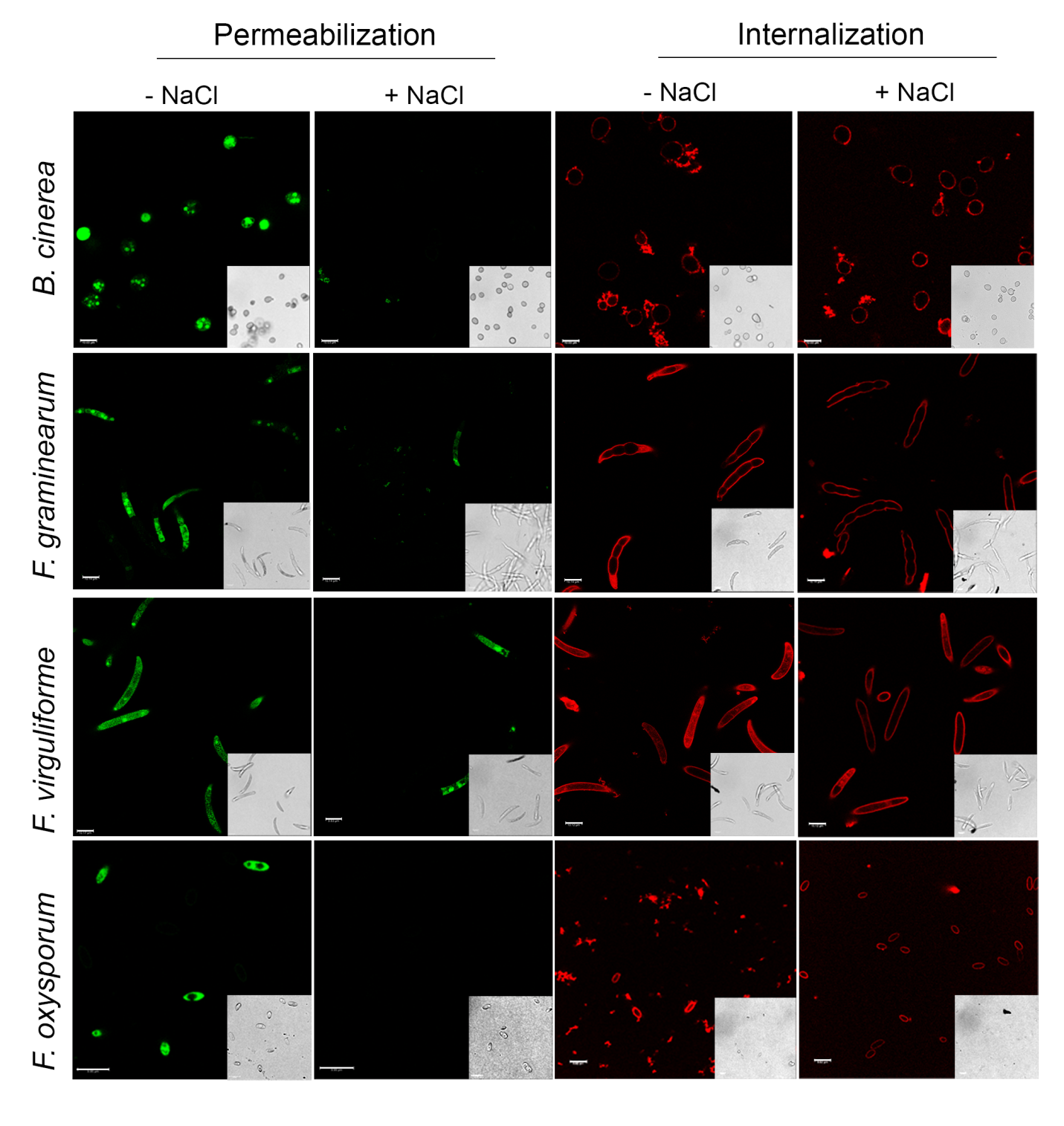

**Fig. S5** Plasma membrane permeabilization of fungal cells and internalization of OefDef1.1 into fungal cells in different pathogens in fungal growth media with or without 100 mM NaCl. Conidia of each fungal pathogen were treated with 2x MIC values of the peptide. Images were captured after 3 h of defensin challenge. SG was used at 1 µM. Bars = 10 μm.

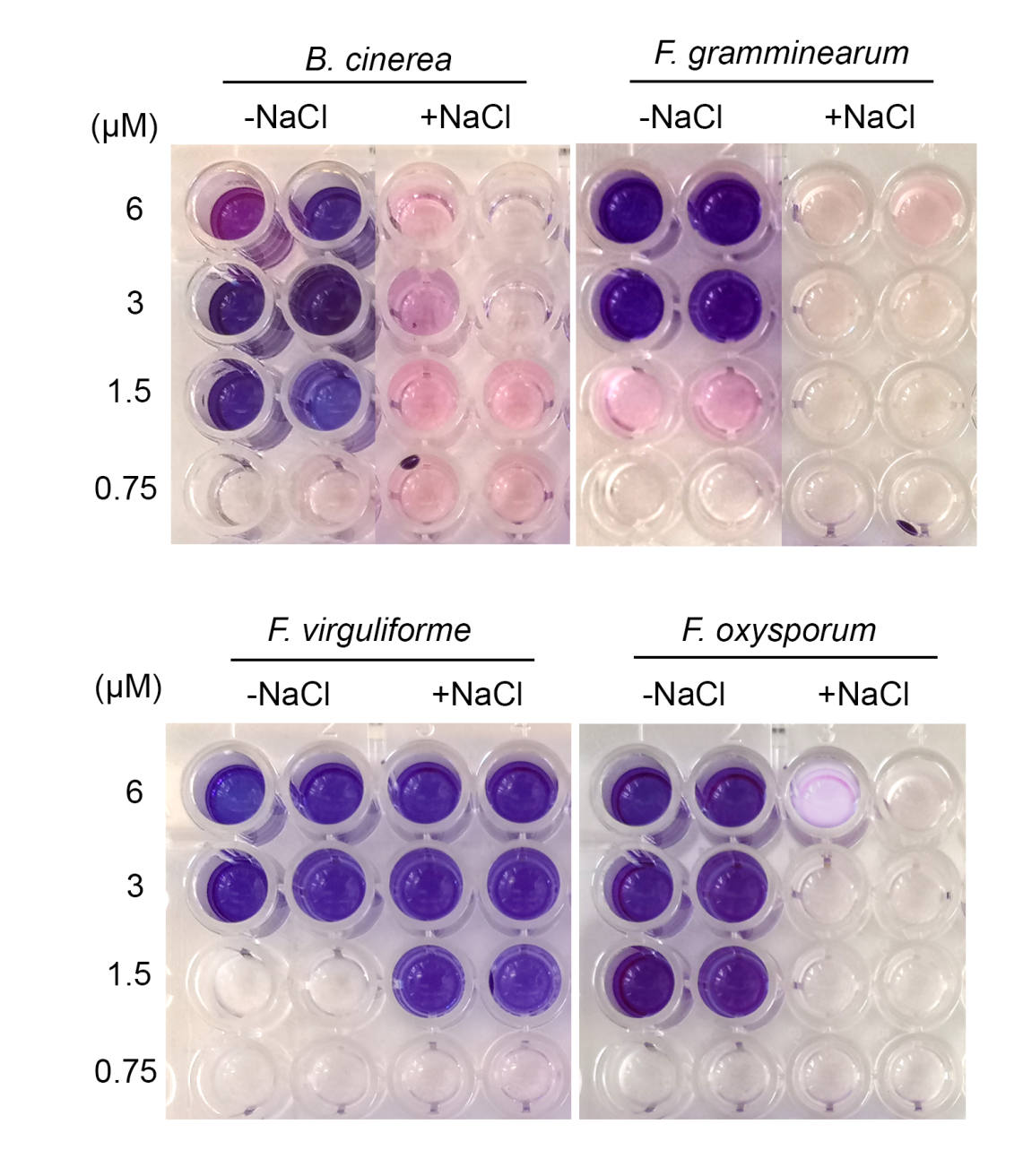

**Fig. S6** MIC value of OefDef1.1 against *B. cinerea*, *F. graminearum*, *F. virguliforme* and *F. oxysporum* in presence of 100 mM NaCl was determined using the resazurin cell viability assay.

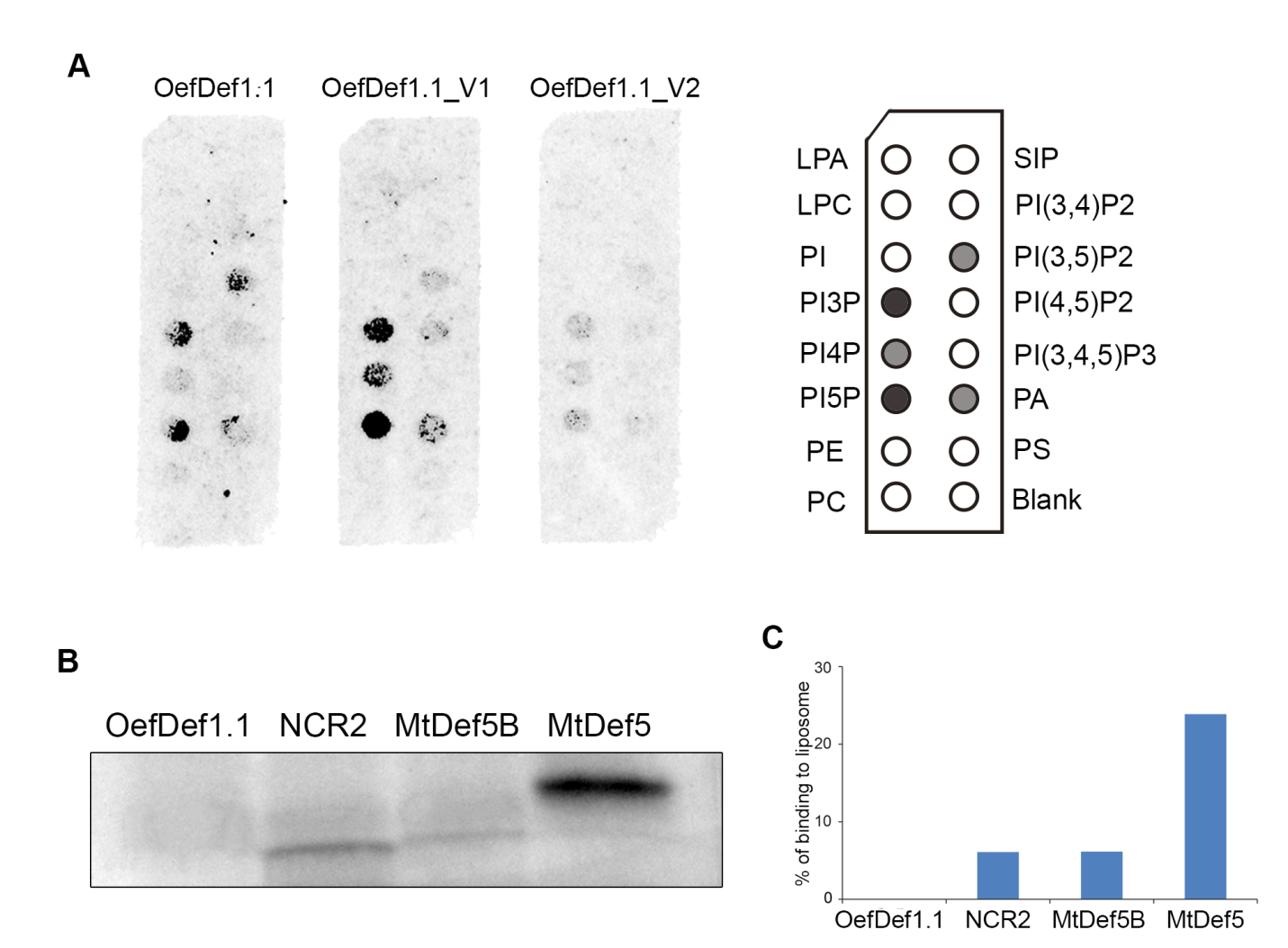

**Fig. S7** Phospholipid binding of OefDef1.1. A. Protein-lipid overlay assay for OefDef1.1, OefDef1.1_V1 and OefDef1.1_V2 using PIP strips. B and C. PI3P liposome binds to NCR2, MtDef5B and MtDef5 peptides (used as positive controls), but not to OefDef1.1. Protein-lipid overlay and PolyPIPosome binding assays were performed as described in Islam *et al.*, 2017.

**Table S1 Characteristics of OefDef1.1 and variants**

| **Peptide** | **Length** | **% hydrophobic amino acids** | **Net charge** | **Molecular Mass**  **(kD)** |
| --- | --- | --- | --- | --- |
| OefDef1.1 | 53 AA | 30% | 9.3 | 6.2 |
| OefDef1.1_V1 | 51 AA | 30% | 10.9 | 5.9 |
| OefDef1.1_V2 | 51 AA | 33% | 8.3 | 5.95 |

**Table S2** Media and conditions for culturing fungal pathogens used in this study

| **Strains** | **Stock medium** | **Spores induced medium** | **Reference** |
| --- | --- | --- | --- |
| *F. graminearum* | PDA | CMC | Cappelini & Peterson (1965) |
| *F. virguliforme* | PDA | PDA |  |
| *F. oxysporum* | PDA | PDA |  |
| *B. cinerea* | PDA | V8 agar | Fermaud & Gaunt (1995) |

CMC indicates carboxymethyl cellulose medium. PDA indicates potato dextrose agar. V8 indicates commercial vegetable juice V8.

**Table S3** Primers used in this study

| **Name** | **Sequences** | **Purpose** |
| --- | --- | --- |
| Bc-ITS-F | TCGAATCTTTGAACGCACATTGCGC | qPCR |
| Bc-ITS-R | TGGCAGAAGCACACCGAGAACCTG |  |
| Lettuce-actin-F | ACATAGCGGGAGCATTGAAC | qPCR |
| Lettuce-actin-R | ACACCCCGTTCTTCTCACAG |  |
| AOX1-F | GACTGGTTCCAATTGACAAGC | Insertion confirmation in pPICZα vector |
| AOX1-R | GCAAATGGCATTCTGACATCC |  |

**Statement of author contributions:**

D. S., S. V. and H. L designed the research. H. L. and S. V. carried out the experiemnts and analyzed the data. H. L. and D. S. wrote the manuscript with input from S. V..
